# A general 3D model for growth dynamics of sensory-growth systems: from plants to robotics

**DOI:** 10.1101/2020.04.21.053033

**Authors:** Amir Porat, Fabio Tedone, Michele Palladino, Pierangelo Marcati, Yasmine Meroz

**Affiliations:** Faculty of Exact Sciences, Raymond and Beverly Sackler School of Physics and Astronomy, Tel Aviv University, Tel Aviv 69978, Israel; Gran Sasso Science Institute, Viale Francesco Crispi 7. 67100, L’Aquila, Italy; Faculty of Life Sciences, School of Plant Sciences and Food Security, Tel Aviv University, Tel Aviv 69978, Israel

**Keywords:** plant tropism, circumnutation, self-growing robots, plant-inspired robotics, 3D, control system, optimal control, growth

## Abstract

In recent years there has been a rise in interest in the development of self-growing robotics inspired by the moving-by-growing paradigm of plants. In particular, climbing plants capitalize on their slender structures to successfully negotiate unstructured environments, while employing a combination of two classes of growth-driven movements: tropic responses, which direct growth in the direction of an external stimulus, and inherent nastic movements, such as periodic circumnutations, which promote exploration. In order to emulate these complex growth dynamics in a 3D environment, a general and rigorous mathematical framework is required. Here we develop a general 3D model for rod-like organs adopting the Frenet-Serret frame, providing a useful framework from the standpoint of robotics control. Differential growth drives the dynamics of the organ, governed by both internal and external cues. We describe the numerical method required to implement this model, and perform numerical simulations of a number of key scenarios, showcasing the applicability of our model. In the case of responses to external stimuli, we consider a distant stimulus (such as sunlight and gravity), a point stimulus (a point light source), and a line stimulus which emulates twining of a climbing plant around a support. We also simulate circumnutations, the response to an internal oscillatory cue, associated with search processes. Lastly we also demonstrate the superposition of both the response to an external stimulus together with circumnutations. Lastly we consider a simple example illustrating the possible use of an optimal control approach in order to recover tropic dynamics, in a way which may be relevant for robotics use. In all, the model presented here is general and robust, paving the way for a deeper understanding of plant response dynamics, as well as novel control systems for newly developed self-growing robots.

## 1 INTRODUCTION

Though the field of robotics has long been inspired from the capabilities of biological organisms, it is only recently that the plant world has become a source of inspiration, particularly due to the ability of plants to continuously change their morphology and functionality by growing, thus adapting to a changing environment (Mazzolai et al., 2016; Mazzolai, 2016; Laschi and Mazzolai, 2016). A new class of plant-inspired robots has emerged, based on the moving-by-growing capabilities of plants. Some recent examples include: (i) a tendril robot developed at NASA’s Johnson Space Center, a slender manipulator composed of multiple bending segments (Mehling et al., 2006), (ii) a vine-bot which elongates its body at the tip by skin eversion, growing along a pre-determined form (Hawkes et al., 2017), and (iii) Plantoid robots inspired by plant roots, based on additive manufacturing technologies (Sadeghi et al., 2014; Dottore et al., 2016; Sadeghi et al., 2017). Though these are impressive accomplishments, these robots are currently limited in their control systems and autonomy. The challenge lies in the fact that the morphology of such self-growing robots changes over time, and is therefore also not known in advance. Furthermore, in the future such robots are expected to perform autonomously in unstructured scenarios, including locomotion in uncertain terrains involving obstacles and voids, as well as the manipulation of unknown objects. Therefore the development of a control system is not trivial, and cannot be based on existing control systems of classic predefined robotic structures.

Plants on the other hand, excel in these type of tasks. Though plants exhibit a variety of types of movements as part of their interaction with their environment (Forterre, 2013), here we focus on the relevant growth-driven movements of rod-like organs such as shoots and roots. Such growth-driven movements are generally classified as either *nastic* or *tropic* (Rivière et al., 2017). Nastic movements are due to internal drivers, such as the inherent periodic movement of plants called circumnutations, sometimes associated with search processes. Tropisms are the growth-driven responses of a plant in the direction of a stimulus, such as a plant shoot growing towards a source of light or away from the direction of gravity (Darwin, 1880; Gilroy and Masson, 2007; Rivière et al., 2017). Tropic responses are based on three main processes: (i) sensing of a directional external stimulus by specialized bio-sensors, (ii) transduction of signals within the plant leading to the redistribution of the growth hormone auxin, resulting in (iii) an anisotropic growth pattern that reorients the organ towards or away from a given stimulus.

In order to emulate these complex growth dynamics in a 3D environment, in a way which is meaningful from the robotics standpoint, a general mathematical framework is required. Recently developed models of growth-driven plant dynamics are limited to specific aspects of tropisms or circumnutations. Bastien et al. have developed models for tropism in 2D, such as the AC (Bastien et al., 2013, 2015) and ACE (Bastien et al., 2014) models, addressing the influence of growth, and identifying the requirement of a *restoring force* called proprioception, whereby a plant can dampen the curving dynamics according to how curved it is (Bastien et al., 2013; Hamant and Moulia, 2016). Bressan et al. (2017) developed a model based on a similar formalism however not accounting for growth explicitly as the driver of dynamics, and achieving stable dynamics by controlling the growth-zone and sensitivity, rather than proprioception. Another model focuses on circumnutations in 3D (Bastien and Meroz, 2016), however disregarding tropic responses.

Here we present a general and rigorous mathematical framework of a rod-like growing organ, whose dynamics are driven both by internal and external cues. Though this model is inspired by plant responses, it is not based on biological details, and is therefore amenable to any rod-like organisms which respond to signals via growth, such as neurons and fungi. The paper is organized as follows: Section 2 describes the dynamical equations of our model based on a 3D description of an organ in the Frenet-Serret formalism, implementing differential growth as the driver of movement, and relating external and internal signals. In Section 3 we present the numerical method required to implement this model, and in Section 4 we perform numerical simulations of a number of key case examples, including responses to external stimuli such as a distant stimulus, a point stimulus, and a line stimulus, as well as circumnutations, the response to an internal oscillatory cue. We also present an example where we superimpose two different types of cues, namely the response to an external stimulus together with circumnutations. Lastly in Section 5 we consider a simple example illustrating the possible use of an optimal control approach in order to recover tropic dynamics, in a way which may be relevant for robotics use.

## 2 GOVERNING EQUATIONS

In this section we develop the dynamical equations at the basis of our model. We first introduce the 3D description of an organ in the Frenet-Serret formalism, followed by the implementation of growth and differential growth as the driver of movement. Finally, we relate external and internal signals to differential growth, which drives the desired movement. We then show that our model is a generalization which consolidates different aspects of existing models, allowing to identify and discuss the characteristic time and length scales of our model.

### 2.1 3D description of an organ

We model an elongated rod-like organ as a curved cylinder with radius *R*, described by its centerline that follows a curve in 3D. We denote the location of the centerline from the origin of a Cartesian frame of reference as 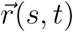, where *t* is time, and *s* is its arc-length, which runs along the organ taking the value *s* = 0 at the base, and *s* = *L* at the apical tip, equal to the total length (see Fig. 11a). In order to describe the dynamics of the centerline with respect to local stimuli, we begin by defining a local frame of reference using the Frenet-Serret framework (Goriely, 2017). Using the Frenet-Serret formulas for a 3D curve parameterization, as shown in Fig. 1a, we can define the tangent vector at arc-length *s* as:

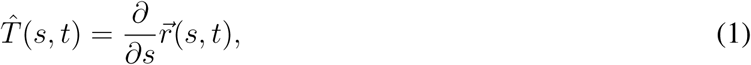

where 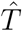 is a unit vector, from the definition of the arc-length (Goriely, 2017). The second derivative of 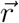 can be written as:

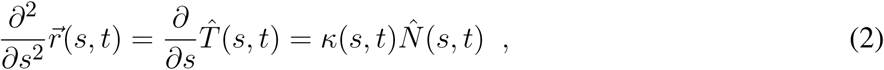

where *κ* is the local curvature of the curve, and 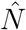 is the respective normal vector. We note that when 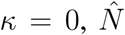 is not defined, in which case we adopt a related local frame described in Section 3. Since 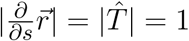, taking the derivative of 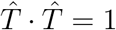 yields 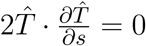, meaning that 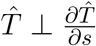, i.e. we have 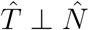. The curvature equals the inverse of the radius of curvature, and the normal vector 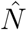 points to the center of the circle with that radius. The third unit vector in the Frenet-Serret framework is the bi-normal vector 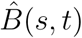, which creates an orthogonal basis in 3D, as illustrated in Fig. 1a:

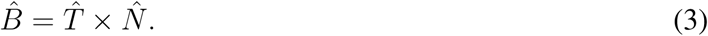

**Figure 1.**
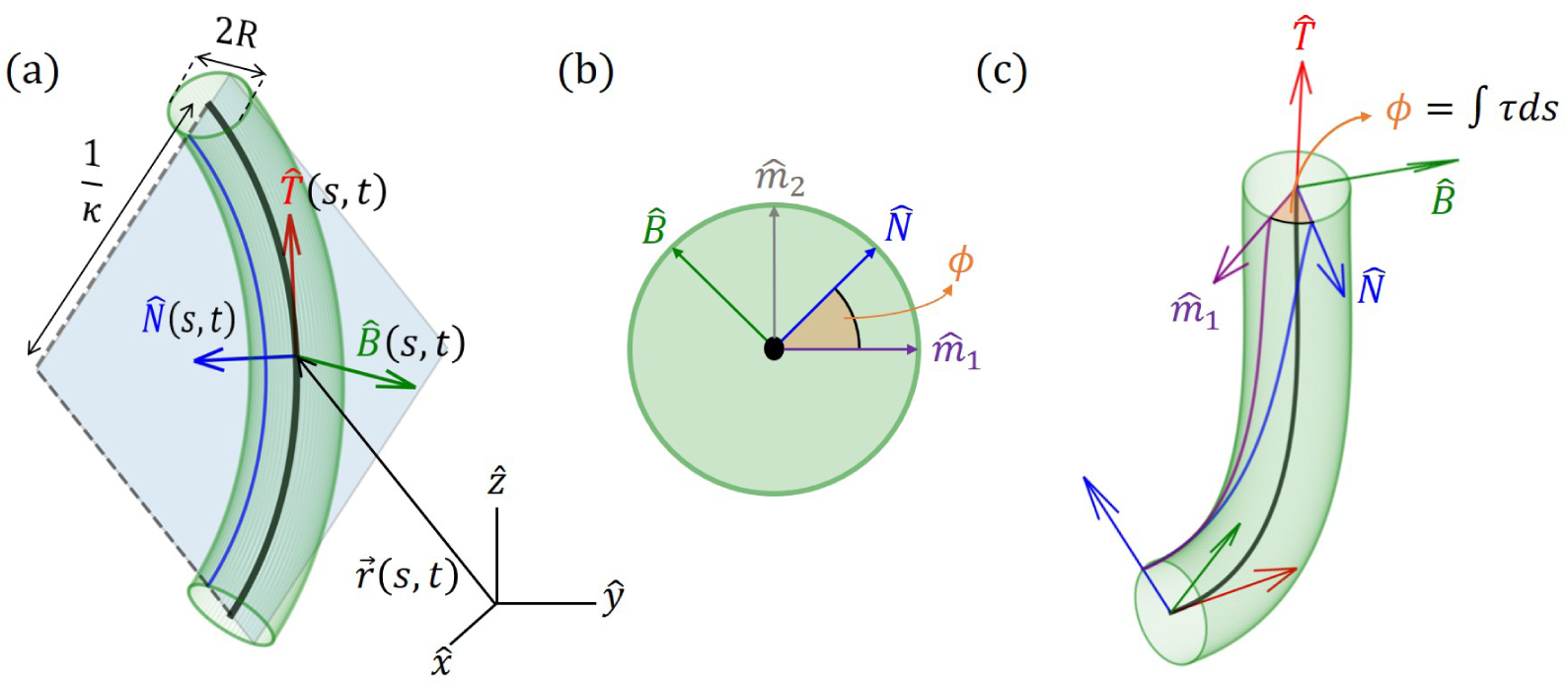
Geometrical definitions for a 3D cylindrical organ. (a) A cylindrical organ of constant radius *R* is described by its centerline, parametrized by the arc-length 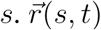 denotes the Cartesian position of a point along the centerline at point *s* and time *t*. The local Frenet-Serret frame at some point along the centerline is defined by the tangent vector 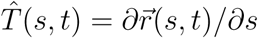, its derivative the normal vector 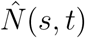 (Eq. 2), and the bi-normal vector 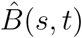 (Eq. 3). Here the organ has a constant curvature *κ* and is restricted to a plane, illustrating 1*/κ*(*s, t*) as the radius of curvature. (b) Cross-section of the organ and the *natural* frame: 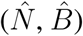 span the cross-section, 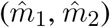 are constant vectors defining the natural frame, as described in Sec. 3, and *ϕ*(*s, t*) defines the angle between 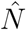 and the reference vector 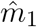. (c) An organ not restricted to a plane. Here *ϕ*(*s, t*) changes along s, and torsion is defined as *τ* = *∂ϕ/∂s*. Note that in (a) *τ* = 0.

For the sake of legibility, we interchangeably omit writing the explicit dependence of variables on (*s, t*), i.e when we write 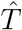 we mean 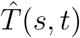.

The Frenet-Serret framework describes the change in this local frame of reference as a function of the arc-length *s* (Goriely, 2017):

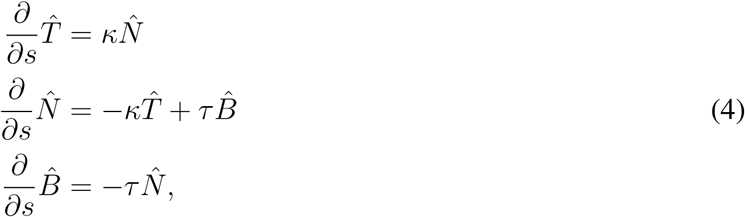

so that the local coordinate system changes accordingly along the curve. Here, *κ*(*s, t*) is the curvature and *τ* (*s, t*) is the torsion of the centerline, describing rotations in the 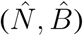 plane leading to a non-planar centerline, as illustrated in Fig. 1b and c. We now define *ϕ*, the angle between 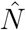 and arbitrarily chosen fixed direction 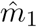 (Bastien and Meroz, 2016; Langer and Singer, 1996; Bishop, 1975). The change in the direction of 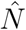 along the curve yields the torsion *τ* (*s, t*) (see Fig. 1c):

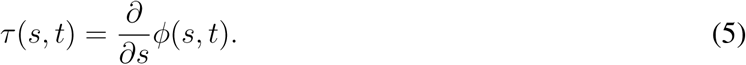

### 2.2 Modeling growth and differential growth

We now introduce growth, using similar definitions to those introduced in (Bastien et al., 2014; Goriely, 2017; Silk, 1989). We define *S*_0_ as the arc-length of the initial centerline of the organ, and the current arc length *s*(*S*_0_, *t*) as the evolution of the point *S*_0_ in time, with initial conditions *s*(*S*_0_, *t* = 0) = *S*_0_.

One can think of the arc-length *s*(*S*_0_, *t*) as describing the flow of the initial point *S*_0_ due to the growth of all previous parts of the organ (see Fig. 2). Therefore, assuming the organ does not shrink, *s*(*S*_0_, *t*) monotonically increases in time. This growth-induced flow within the organ motivates us to use definitions from fluid dynamics, in which the parameter *S*_0_ can be thought of as the Lagrangian, referential, or material coordinate, and *s* as the Eulerian or spatial coordinate (Goriely, 2017). Using regular conventions of continuum mechanics, we define the local velocity of point *s* as the accumulation of the growth that occurs in previous parts of the organ:

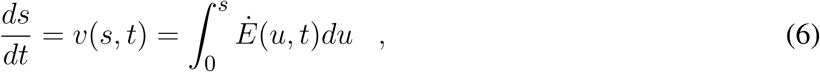

where *Ė*(*s, t*) is the local growth rate, representing a combination of the effect of addition of new cells and their elongation. We define the length of the active growth-zone of a growing organ *L*_*gz*_ as the length over which the growth rate *Ė*(*s, t*) is non-zero. In plants growth is generally confined to a finite sub-apical growth-zone: *L* − *L*_gz_ ≤ *s* ≤ *L*, as shown in Fig. 2. We note that as opposed to Lagrangian quantities (functions of *S*_0_), the time derivative of Eulerian fields (functions of *s*(*S*_0_, *t*)) incurs an additional convection term. We use the convention of material derivatives for the total time derivative, namely: *D/Dt* ≡ *∂/∂t* + *v∂/∂s*.

**Figure 2.**
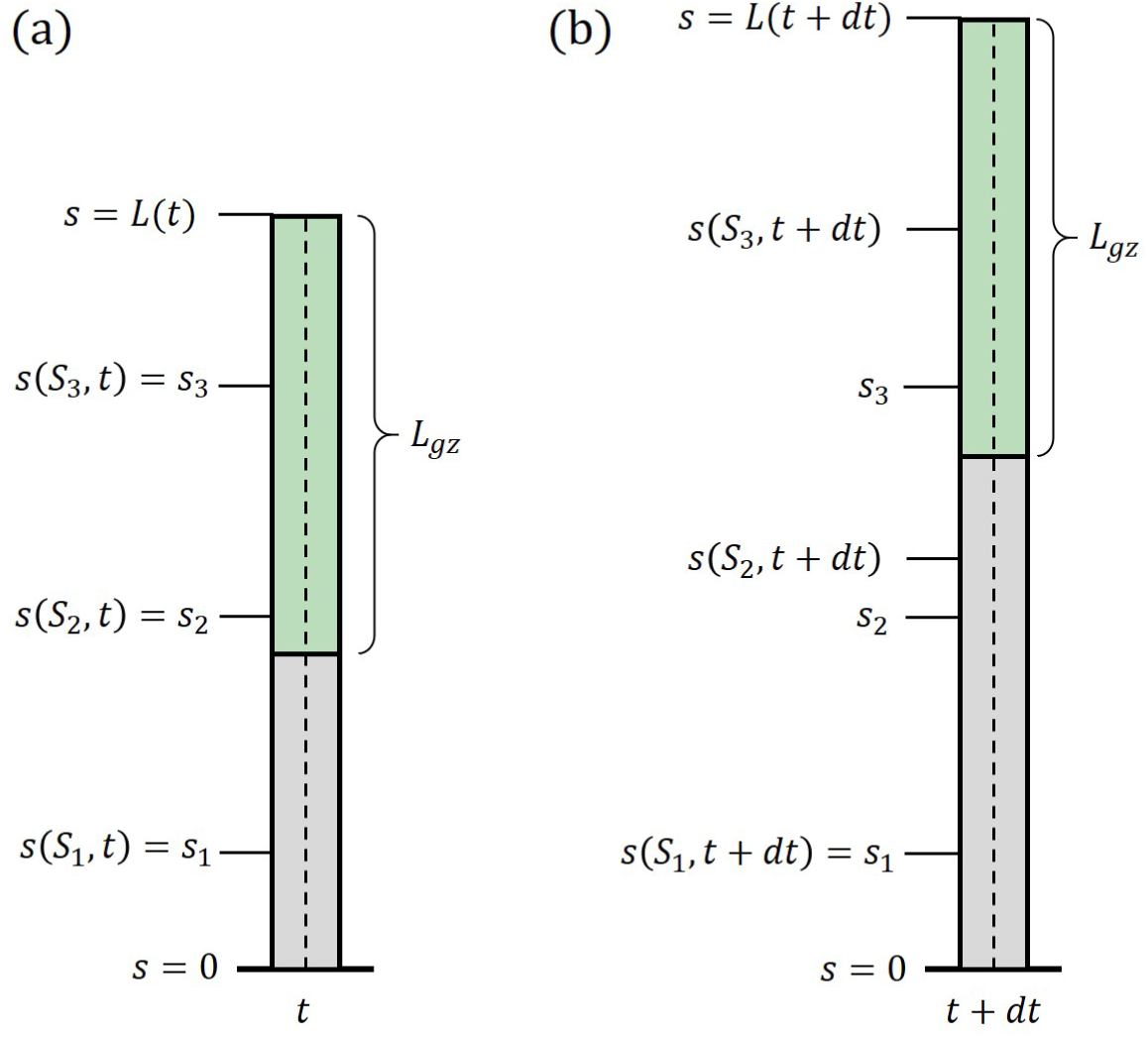
Growth description. Illustration of a growing organ with a sub-apical growth zone, marked in green. The centerline (dashed line) can be parameterized by a material coordinate, *S*_0_, or by the arc-length, *s*(*S*_0_, *t*). (a) the organ at time *t*, (b) the organ at time *t* + *dt*. Outside of the growth zone, the position of the material coordinate does not change in time *s*(*S*_1_, *t* + *dt*) = *s*(*S*_1_, *t*). Within the growth zone *s*(*S*_3_, *t* + *dt*) *> s*(*S*_3_, *t*), i.e. the location of the material coordinated flows due to growth, and *S*_2_ and *S*_3_ flow within the organ. *S*_2_ flows out of the growth zone and will stay fixed.

As mentioned in the Introduction, plant tropisms are the growth-driven reorientation of plant organs due to a directional stimulus such as light, gravity, water gradients etc. In particular, the reorientation of the plant organ is due to differential growth, i.e. one side of the cylindrical organ grows at a higher rate than the other side, resulting in a curved organ. Following Bastien and Meroz (2016), we consider an infinitesimal cross-section of a cylindrical organ, and define the differential growth rate in a direction *ê* as the difference in growth rate 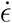 on either side, normalized by their sum:

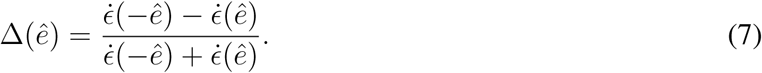

Following this definition, for Δ(*ê*) > 0 the organ grows faster in the direction –*ê* and the organ therefor bends in the the direction *ê* (see Fig. 3). We now define the differential growth vector, which is in the direction of the active reorientation:

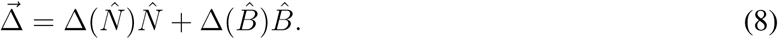

**Figure 3.**
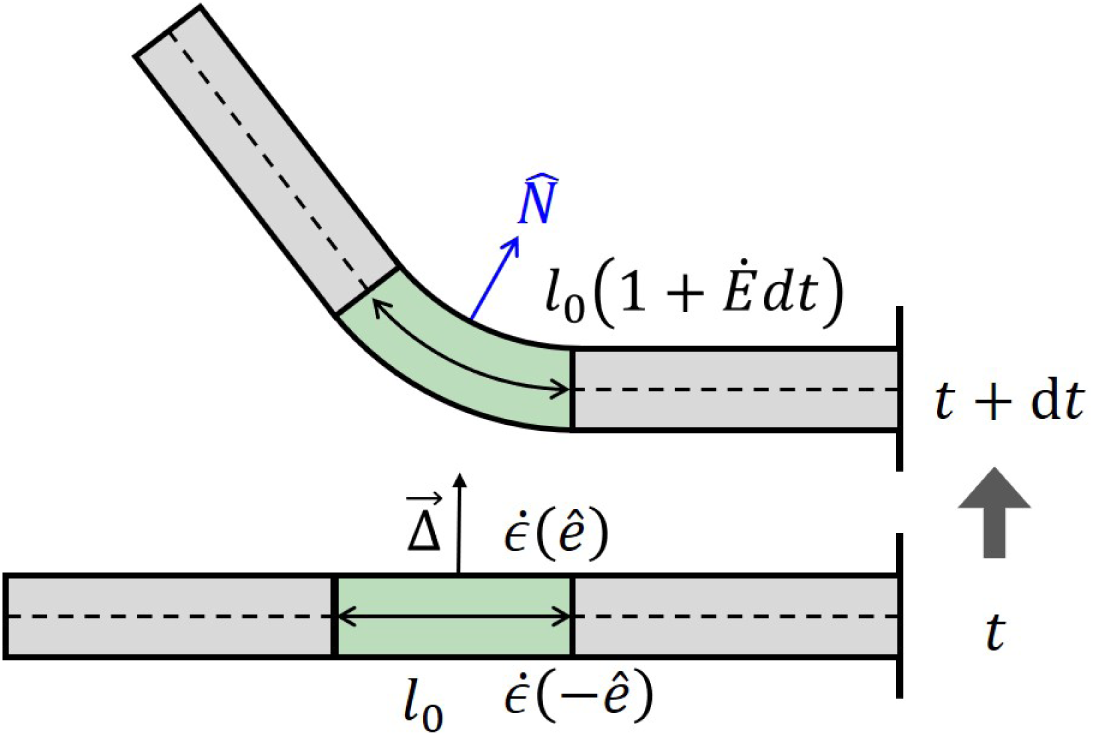
Differential growth. Differences in growth rates across a cylinder, lead to a change in curvature. At time *t* we have a straight organ with *κ*(*t*) = 0, and with a growth zone in the center of length *l*(*t*) = *l*_0_, marked in green. The differential growth vector 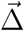 in the growth zone is constant and points upwards in the *ê* direction. Following Eq. 7 the growth rate on the lower side is higher than in the upper side 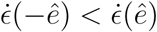, and after a time interval *dt* the two sides grow different amounts, leading to bending of the growth zone with a new curvature *κ*(*t* + *dt*) *>* 0. The new length of the growth zone along the centerline is now *l*(*t* + *dt*) = *l*_0_(1 + *Ėdt*). Note that changes in curvature in the middle of the organ lead to changes in orientation of the rest of the organ.

In order to describe the active reorientation of an entire organ, we relate the shape of the organ and its growth dynamics, expressed by the dynamics of its local curvature, 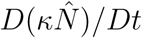, to the differential growth term 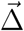 (Bastien and Meroz, 2016), *resulting in:*

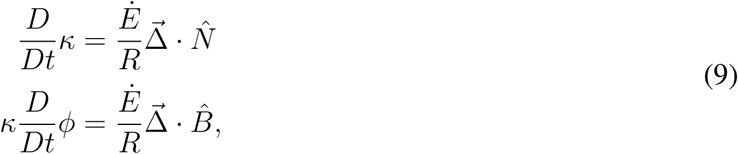

where the equations have been linearized by assuming that the radius of curvature 1*/κ* is always larger than the radius of the organ (*κR* ≪ 1). For a detailed calculation see Appendix A in the Supplementary Material (SM). These equations are similar to those developed in (Bastien and Meroz, 2016), where the differential growth vector represented the internal cues related to circumnutations. Given an expression for the differential growth vector 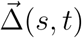 and an initial configuration, the dynamics can be integrated completely. The form of 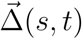 is dictated by either internal cues (circumnutations) or external stimuli, as discussed in the following section.

### 2.3 Relating external and internal signals to differential growth

In the last section we represented the anisotorpic growth pattern by the local growth differential growth vector 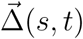. An external signal is translated to a specific growth pattern thanks to signal-specific bio-sensors and biochemical signal transduction mechanisms. Here we reduce these complex processes to a sensitivity or gain function, which maps the external signal to a growth response, and directly governs the growth differential vector.

Environmental signals can be mathematically described as fields. For example vector fields describe light and gravity, while a scalar field describes the concentration of water or nutrients, and the direction of increasing concentrations is again described by a vector field of the gradients. Lastly tensor fields may describe stress and strain, however we will not discuss these here since our model does not include elasticity. Here we focus on vector fields, where we can write the directional stimulus in the form 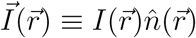, where 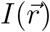 is the magnitude of the stimulus at point in space, and 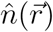 is its direction. For example, in the case of an infinitely distant stimulus such as light and gravity, the stimulus magnitude and direction is constant in space, i.e. 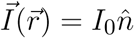. In the case of a chemical concentration gradient, a possible form would be 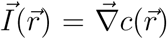, though the sensed magnitude may depend on other factors such as the concentration itself, and remains to be verified. The physics of the signal and the geometry of the emitting source dictate the direction of the stimulus 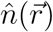. Within a specific infinitesimal element of an organ, the differential growth vector is restricted to the cross-section, i.e. the local 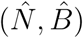 plane. Therefore the relevant directional information of the stimulus lies within its projection perpendicular to the organ surface, as illustrated in Fig. 4. We define the component of the stimulus perpendicular to this surface, 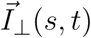, as the effective stimulus sensed by the organ. From geometrical arguments, the effective stimulus sensed by a cylindrical surface is given by:

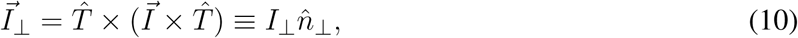

where we have defined 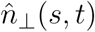 as the direction of the perpendicular component of the signal, and *I*_⊥_(*s, t*) its magnitude given by *I*_⊥_ = *I*_0_ sin (*θ*(*s, t*)), where *θ*(*s, t*) is the angle between the surface 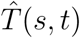 and the direction of the stimulus 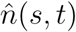, as shown in Fig. 4.

**Figure 4.**
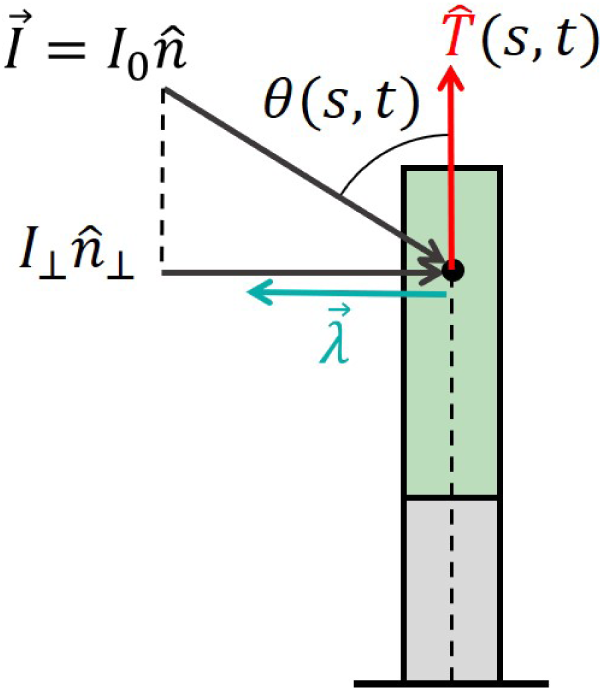
Effective signal and response vector. An example of a signal that can be described by a constant vector field (such as sunlight and gravity) of the form 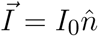, where *I*_0_ is the magnitude of the stimulus, and 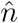 is its direction. For an element of an organ, 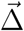 the relevant directional information of the stimulus lies within its projection perpendicular to the organ surface, which we define as 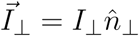 (see Eq. 10), where the magnitude is given by *I*_⊥_ = *I*_0_ sin (*θ*(*s, t*)), and *θ*(*s, t*) is the angle between the surface 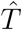 and 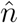. Biophysical laws generally describe sensory responses to input signals as functions of the signal intensity *λ*(*I*) (see main text). We define the response vector 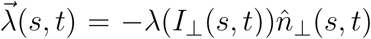 where the magnitude of the perceived response is given by the sensitivity function *λ*(*I*_⊥_ (*s, t*)), and 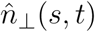 is the direction of the effective stimulus.

Two central biophysical laws describe sensory responses to input signals, which we term here the sensitivity function *λ*(*I*). One is a logarithmic relationship *λ*(*I*) = *a* + *b* log (*I/I*_0_), referred to as the Weber-Fechner law (Norwich and Wong, 1997), and the other is a power law relationship *λ*(*I*) = *aI*^*b*^, known as Stevens law (Stevens, 1957). As an example it has been found that phototropism follows Stevens’ Law (Bastien et al., 2015), while in the case of gravitropism only inclination is sensed, and sensitivity is constant, *λ*(*g*) = const (Chauvet et al., 2016). However very little is known for other plant tropisms. We now define the local response vector:

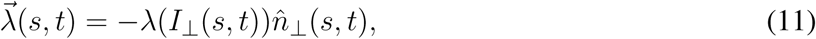

where the sensitivity function takes the effective stimulus sensed by the organ *λ*(*I*_⊥_(*s, t*)), and 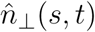 is the direction of the effective stimulus. As stated before, the differential growth vector is restricted to the cross-section plane of the organ element, and it is therefore directly related to the perpendicular component of the intensity, and its response vector, i.e. 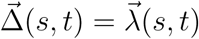.

However, it has been found that a so called *restoring force* is required for stable posture control, termed *proprioception* (Bastien et al., 2013; Hamant and Moulia, 2016). This is related to an internal process associated with the active tendency of a growing organ to resist being bent (not a mechanical response), and is represented by 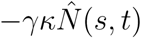, where *γ* is the proprioceptive sensitivity (Bastien et al., 2013). We also note that differential growth may be due to internal processes such as in the case of circumnutations. Here an internal oscillator turns the differential growth vector in the 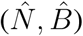 plane (Bastien and Meroz, 2016), and can be described as 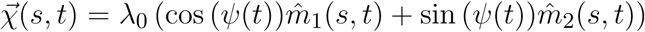, where *λ*_0_ is the intensity of the bending, and *ψ*(*t*) is a general function describing the direction of growth at time *t*, relative to fixed vectors 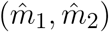 (Fig. 1b). Here we chose a circular growth pattern, however more elaborate forms can be implemented (Bastien and Meroz, 2016). Adding the propriocetion term and circumnutations, the differential growth vector therefore follows:

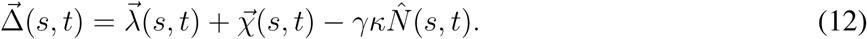

Together with with Eqs. 9, Eq. 12 completes our model for active growth-driven movements of rod-like organs in 3D, taking into account external signals, internal cues (circumnutations), and posture control. For multiple stimuli, assuming additivity one can replace 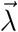 with the sum of specific response vectors 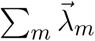 (Bastien et al., 2015). A number of specific cases, including various types of external and internal cues, are explained in further detail in Section 4. A schematic summarizing the governing equations is brought in Fig. 5.

**Figure 5.**
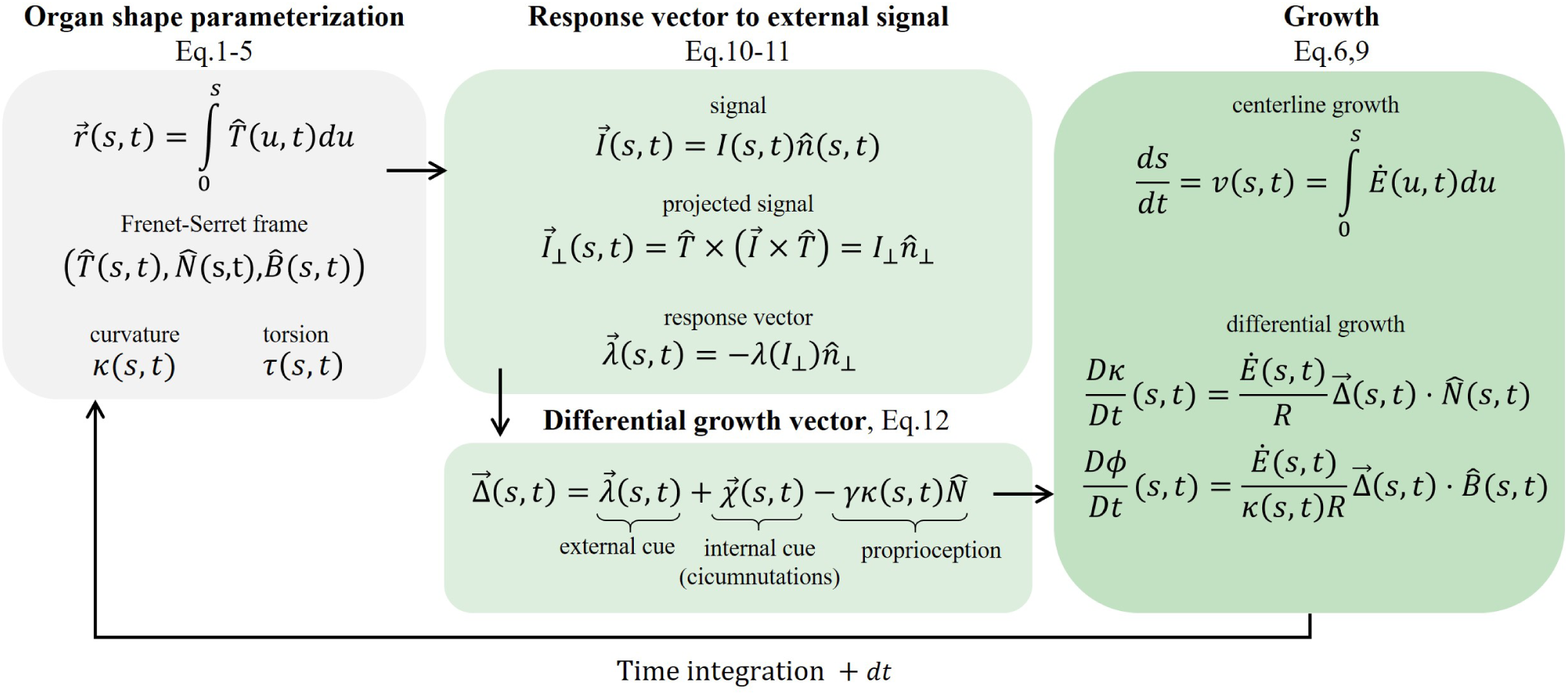
Schematic of the governing equations. We present the main stages involved in the model: (a) Organ shape parametrization, Section 2.1: the Frenet Serret local frame in Eqs. 1-3, and Frenet-Serret equations which also define *κ* and *τ*, brought in Eq. 4. (b) Response vector to external signal, Section 2.3: assuming a vector field signal, we find the projected signal (Eq. 10), and calculate the response vector (Eq. 11) which affects the growth response. (c) Differential growth vector (Eq. 12), includes terms representing external cues (the response vector), internal cues (circumnutations), and proprioception for posture control. (d) Implement growth dynamics, Section 2.2. The centerline is updated using Eq. 6 and Eq. 9, using the constructed differential growth vector.

Lastly, the distribution of sensory systems along the organ also requires attention. Sensory systems in plant organs are generally either distributed along the organ, providing local sensing (Sakamoto and Briggs, 2002; Wan et al., 2008; Hohm et al., 2013), or restricted to the tip, termed apical sensing (Darwin, 1880; Holland et al., 2009; Hohm et al., 2013; Knieb et al., 2004). For example, in the case of shoot phototropism, photoreceptors are localized at the tip alone, such as in wheat, or distributed along the whole growth zone as in the case of Arabidopsis. In the case of gravitropism, specialized cells called statocytes sense the direction of gravity, and these are generally found throughout the growth zone for aerial organs, and restricted to the tip for roots (Morita and Tasaka, 2004; Su et al., 2017). In the case of apical sensing, the local response vector 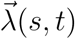 will be replaced with that of the apex 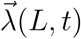, meaning that the whole organ responds to what is sensed at the tip alone.

### 2.4 Comparison to previous models

Different models of growth-driven plant dynamics have been recently developed, encompassing different aspects of tropisms and circumnutations. Bastien et al. (Bastien et al., 2013, 2014, 2015) have developed a model for tropism in 2D, addressing the influence of growth, and identifying the requirement of proprioception. A third model (Bressan et al., 2017) is based on a similar formalism, achieving stable dynamics by controlling the growth-zone and sensitivity, rather than proprioception. Another model focuses on circumnutations in 3D (Bastien and Meroz, 2016), disregarding tropic responses. In what follows we show how our model relates to these previous models, while also generalizing them and unifying them.

In order to compare with 2D models, we focus on the case where the dynamics of our model are restricted to a 2D plane, which occurs when the direction of the stimulus 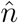 is in the plane defined by 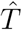 and 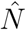. In this case, following Eq. 12, only the component in the 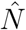 direction of 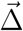 is not zero. Substituting this in the dynamical equations in Eq. 9, since 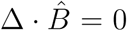 we get that *∂ϕ/∂t* = 0, i.e. *ϕ* is constant. Assuming an initially straight organ, *ϕ* = 0 throughout, yielding *τ* = *∂ϕ/∂s* = 0. The geometrical meaning is that when the stimulus and the initial state of the organ are in the same 2D plane, the dynamics of the organ will remain within that plane, and therefore restricted to 2D. In this case we can compare the dynamics directly to (Bastien et al., 2015) by projecting the model to 2D, and assuming a constant signal. We define the local angle of 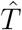 along the organ with respect to the direction of the constant signal 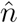, as illustrated in Fig. 4, i.e. 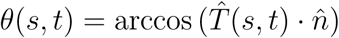. Taking the derivative over the arc-length *s*, and recalling that 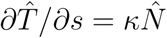 (Eq. 4) yields: 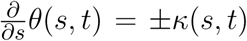, where the sign depends on the direction of 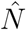. Substituting these expressions in Eqs. 12 and 9, together with 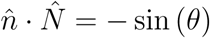 and a constant sensitivity function *λ*(*I*) = *λ*, we get:

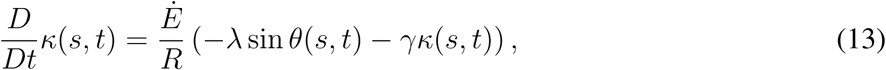

identical to the ACE model developed in (Bastien et al., 2014).

We now consider (Bressan et al., 2017). Their main equation of motion appears in Eq. (2.8), and translating this into our terminology takes the form:

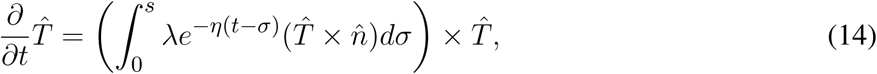

where *λ* > 0 is a constant measuring the strength of the response, similar to our tropic sensitivity, while *e*^−*η*(*t*−*σ*)^ is what they call a stiffness factor. The simplest way to compare this model is by looking at its 2D projection. Taking 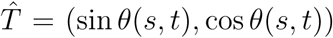, where *θ*(*s, t*) is the angle between 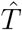 and 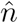, and substituting this in Eq. 14 leads to 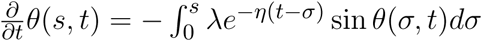. Taking a derivative in s finally yields:

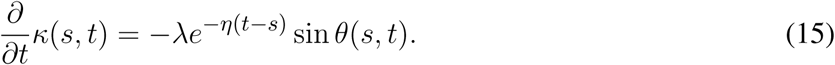

We note that this model considers accretive growth, where material is added at the tip, and elongation is disregarded. This means that growth is only taken into account implicitly as the driver of the tropic movement, and a material derivative is not required, a good approximation of the dynamics in certain cases (Bastien et al., 2013, 2014). In this case the ACE model in Eq. 13 converts to the AC model:

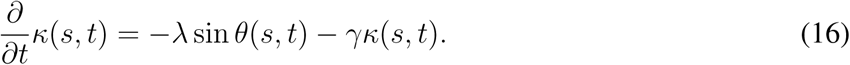

Comparing Eqs. 15 and 16, we see that the equations are similar: the response, appearing in the l.h.s., is identical, and in the r.h.s. the tropic stimulus is represented by sin *θ*(*s, t*) in both, as well as a sensitivity factor. In Bressan’s model the stiffness prefactor *e*^−*η*(*t*−*s*)^ represents a smooth growth zone with a characteristic size of 1*/η*: the youngest parts (s=t at the tip) the stiffness factor is 1, while older parts of the organ (as s goes to zero) the stiffness factor goes to 0. We also notice that Bressan do not use a proprioceptive term, generally required for stable dynamics, however they were able to circumvent this problem by using small growth zones.

### 2.5 Characteristic length and time scales

In Section 2.4 we show that in the case where the dynamics of our model are restricted to a 2D plane our model recovers the ACE model developed by (Bastien et al., 2014). Thanks to this relation, we can adopt their dimensional analysis (Bastien et al., 2013) which identifies characteristic length and time scales. Consider the case of a constant stimulus placed perpendicular to a shoot. The length scale is identified by considering the steady state, where the shoot has grown in the direction of the stimulus achieving a steady state form, with 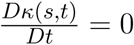 everywhere, including the growth zone. Substituting this in Eq. 13 yields the maximal curvature value *κ*_max_, and its inverse, the radius of curvature, corresponds to a characteristic length scale termed the *convergence length L*_*c*_ = 1*/κ*_max_ = *γ/λ*, where *γ* and *λ* are the proprioceptive and tropic sensitivities respectively. There are two time scales: one is associated with the time it takes for the organ to reach its steady state, termed the *convergence time* and defined as *T*_*c*_ = *R/Ėγ*. The other is associated with the time it takes the organ to align in the direction of the stimulus for the first time, termed the *arrival time*, as is defined as *T*_*v*_ = *R/ĖL*_gz_*λ*. The ratio between the convergence length *L*_*c*_ and the length of the growth zone *L*_gz_, as well as the ratio between the convergence time *T*_*c*_ and arrival time *T*_*v*_, introduces a dimensionless number *B*, termed the *balance number* (Bastien et al., 2013; Hamant and Moulia, 2016), which describes the balance between the sensitivity to external stimuli and proprioception, and is linearly related to the maximal curvature:

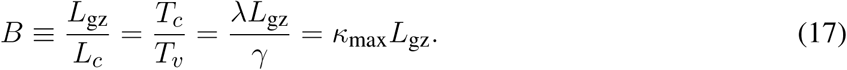

Low values of *B* mean that *L*_*c*_ > *L*_gz_, i.e. the growth zone is not big enough to contain the full bending within the given curvature, or alternatively that *T*_*v*_ > *T*_*c*_, i.e. the organ dynamics converge before it is able to arrive to the desired orientation in the direction of the stimulus. High values of *B* mean that *L*_*c*_ < *L*_gz_, i.e. the growth zone can contain the full bending, or alternatively that *T*_*v*_ < *T*_*c*_, i.e. the organ arrives to the desired orientation before the dynamics converge, therefore also exhibiting damped oscillations. In other words we see that the balance number *B* represents a relation between the final shape of the organ in steady state, and the dynamics.

## 3 NUMERICAL METHOD

### 3.1 Natural frame for numerical scheme

As stated in Section 2, our model for active growth-driven dynamics, described by Eqs. 9 and 12 and schematically illustrated in Fig. 5, is formulated in the Frenet-Serret frame. The Frenet-Serret frame is a natural choice to describe curves since the second derivative gives the local curvature, 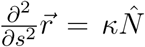, a natural geometrical quantity. However within this framework 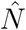 is not defined when *κ* = 0. In order to avoid related numerical issues, in the numerical scheme we adopt a related local frame termed the “natural frame” (Bishop, 1975; Langer and Singer, 1996): assuming 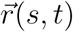 is a point along the centerline of the organ, the natural frame is described by the orthonormal vectors 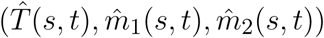, where 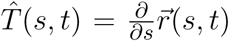 is the tangent vector in Eq. 1. The other two orthogonal vectors 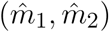 span the cross section plane spanned by 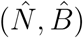 in the Frenet Serret frame. The rotations of this local frame with respect to the arc length of the curve is described using the following equations, similar to the Frenet-Serret equations in (Eq. 4):

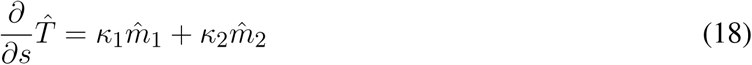

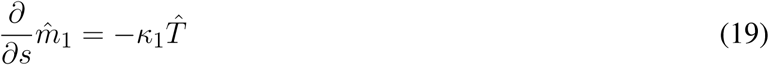

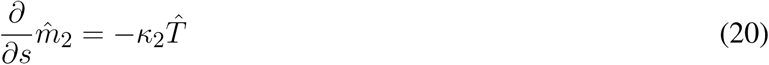

Here, *κ*_1_(*s, t*) and *κ*_2_(*s, t*) are the curvature components of the local cross section plane, and the total curvature *κ*(*s, t*) and torsion *τ* (*s, t*) are given by the relations:

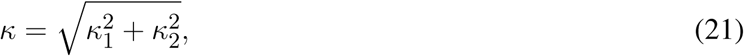

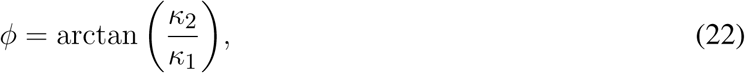

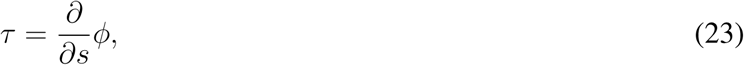

where *ϕ* is the angle between 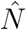 (in the Frenet-Serret frame) and 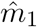, illustrated in Fig. 1b, used to define *τ* in Eq. 5. This frame is closely related to the Frenet-Serret frame, however the cross-section directions 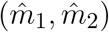 are always well defined, even when *κ* = 0. Within this frame, Eqs. 9 can be rewritten as (see Appendix A in the SM for a detailed calculation):

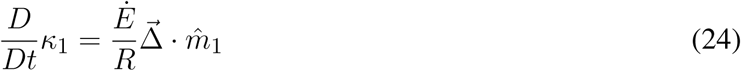

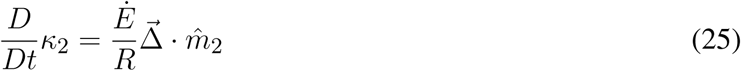

In order to solve the dynamics we integrate Eqs. 24-25.

### 3.2 Discretization and integration

The organ is divided into segments of length *ds*, and we rewrite functions of the centerline in a discrete form, following the general form:

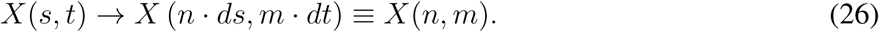

We describe the location of the organ using the local coordinate system:

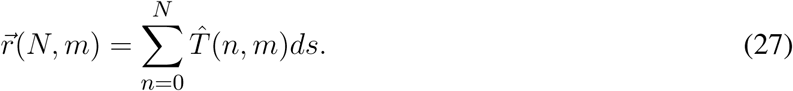

The dynamics of the organ is described through the evolution of the local coordinate system. We rewrite Eq. 18-20 in matrix form, which describe the change in the local frame of reference as a function of *s*:

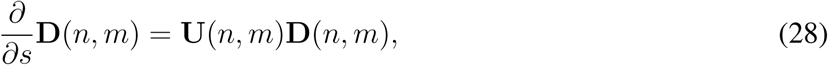

where **D**(*n, m*) is the rotation matrix:

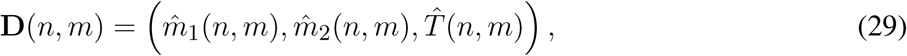

and **U**(*n, m*) is the skew symmetric Darboux matrix:

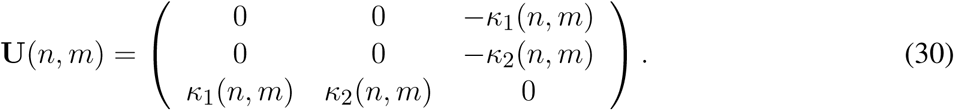

In order to integrate Eq. 28 we use the Rodrigues formula (Gazzola and L, 2018):

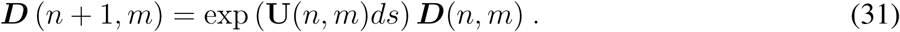

It is therefore enough to find the evolution of **U**, or the evolution of *κ*_1_ and *κ*_2_, to describe the organ in time. To integrate *κ*_1_ and *κ*_2_, we discretize Eqs. 24 and 25, adopting the following numerical time and arc-length derivatives (where *dt* is the discretized time step):

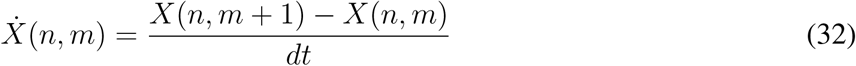

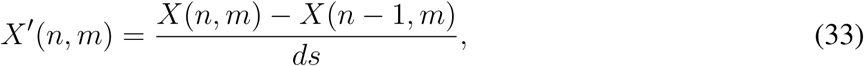

leading to:

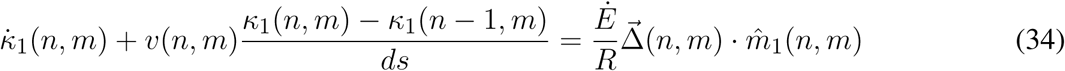

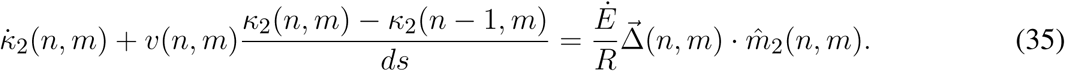

The growth speed appearing in the material derivative, *v*(*n, m*), is calculated following Eq. 6. Assuming a growth-zone of length *L*_gz_ and uniform growth rate *Ė*, leads to:

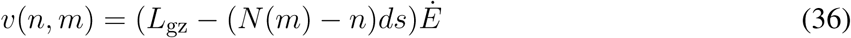

in the case *L*_gz_ ≥ (*N* (m) − *n*)*ds*, and *v*(*n, m*) = 0 otherwise. Extracting 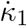 and 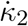 from Eqs. 34-35, we substitute these in Eq. 32. Together with the following straight initial conditions and clamped boundary conditions of the organ, we integrate over time:

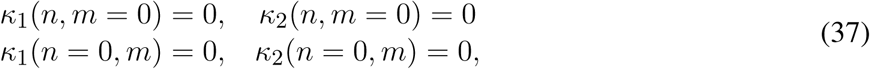

finally resulting in *κ*_1_(*n, m* + 1) and *κ*_2_(*n, m* + 1). In order to find the proper relation between spatial and temporal discretization, we consider the equation for the velocity at the tip, in Eq. 36, in which case *N*(*m*) = *n*, and recalling that 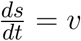 yields the relation:

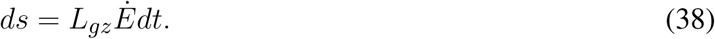

### 3.3 Implementing growth

As discussed in Section 2.2, growth is implemented via a material derivative with a local growth rate described in Eq. 6, representing the elongation of cells in the growth zone, creating a one dimensional growth flow within the organ. When cells reach a certain threshold size they stop elongating, thus leaving the size of the growth zone *L*_gz_ constant. Since the total length of the organ increases with time, in the numerical scheme we add a new segment *ds* at the tip at each time step:

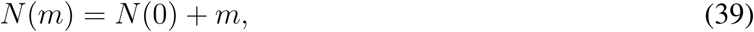

where N(m) is the total number of segments in the organ at time step m, and therefore the total length is *L*(*m*) = *N*(*m*) · *ds*. This is not to be confused with accretive growth where material is added at the tip alone. Special care is required in assigning the correct curvature values to the newly added segments. At time *m* − 1 we initialize the next *N*(*m*)-th segment so that *κ*_1_(*N, m* − 1) = 0, *κ*_2_(*N, m* − 1) = 0, 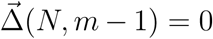, and *v*(*N, m* − 1) = *L*_*gz*_*Ė* (the velocity at the tip as defined in Eq. 6). Substituting these values in Eq. 34-35 yields *κ*_1_ (*N, m*) = *κ*_1_ (*N* − 1, *m* − 1) and *κ*_2_ (*N, m*) = *κ*_2_ (*N* − 1, *m* − 1), i.e. the curvature of the new segment is identical to the its predecessor.

### 3.4 Simulation parameters

In the simulations presented in the next section, the initial conditions include a straight vertical organ *κ*(*s, t* = 0) = 0 (i.e *κ*_1_(*s, t* = 0) = 0 and *κ*_2_(*s, t* = 0) = 0), with an initial length *L*_0_ = 1.0 and a growth zone *L*_*gz*_ = 1.0. Boundary conditions are defined with a clamped base *κ*(*s* = 0, *t*) = 0 (*κ*_1_(*s* = 0, *t*) = *κ*_2_(*s* = 0, *t*) = 0). The organ radius is *R* = 0.1, the proprioceptive coefficient is *γ* = 0.01, and the tropic sensitivity (when applicable) was taken to be either *λ*_0_ = 0.1 or *λ*_1_ = 0.05. The ratio of the proprioceptive and tropic sensitivity values substituted in Eq. 17, correspond to balance numbers *B* = 10 and *B* = 5 accordingly, both in the range of what has been observed in plants (Bastien et al., 2013). The maximal curvature is *κ*_max_ = *λ*_0_*/γ* = 10, yielding *κ*_max_*R* = 1. This means that *κR* ≤ 1 throughout the simulations, in agreement with the low curvature assumption. The simulation time step is *dt* = 0.1, and the length of the discrete elements is *ds* = 0.01. A constant growth rate was taken all along the growth zone following Eq. 38: *Ė*= *ds/dtL*_*gz*_ = 0.1. In the next section we discuss simulations of specific cases. The code is freely available at https://github.com/poratamir/3D-growth-dynamics.

## 4 CASE EXAMPLES AND SIMULATIONS

Here we discuss various representative cases of internal and external cues. Since the differential growth term is the driver of the dynamics, it is the only term which needs to be defined accordingly. We bring the specific form of the differential growth vector for each case, as well as a snapshot of a numerical simulation. Videos of the full simulation dynamics can be found in the SM, also showing the evolution of *κ* and *τ* over time.

### Infinitely distant constant stimulus

The simplest type of stimulus is a constant stimulus placed at infinity. In this case the stimulus is a parallel vector field originating from direction 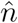, and constant in space and time, i.e. 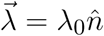:

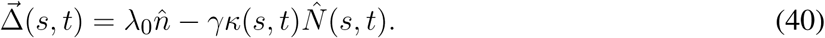

The sensitivity *λ*_0_ may depend on the intensity of the stimulus, for example in the case of phototropism, following either the Weber Fechner or Stevens’ Law, as discussed in Section 2.3. This is not the case for gravitropism, since plants sense inclination rather than acceleration (Chauvet et al., 2016). A snapshot of the final form of the simulation is shown in Fig. 6a, and and example of the full dynamics can be found in Video 1 in the SM. Since the projection of this equation in 2D yields the ACE model (Bastien et al., 2013, 2014), we validate our model numerically, showing that our 3D simulations converge to the known analytical solution in 2D with an exponential growth profile (see Appendix B in the SM for details).

**Figure 6.**
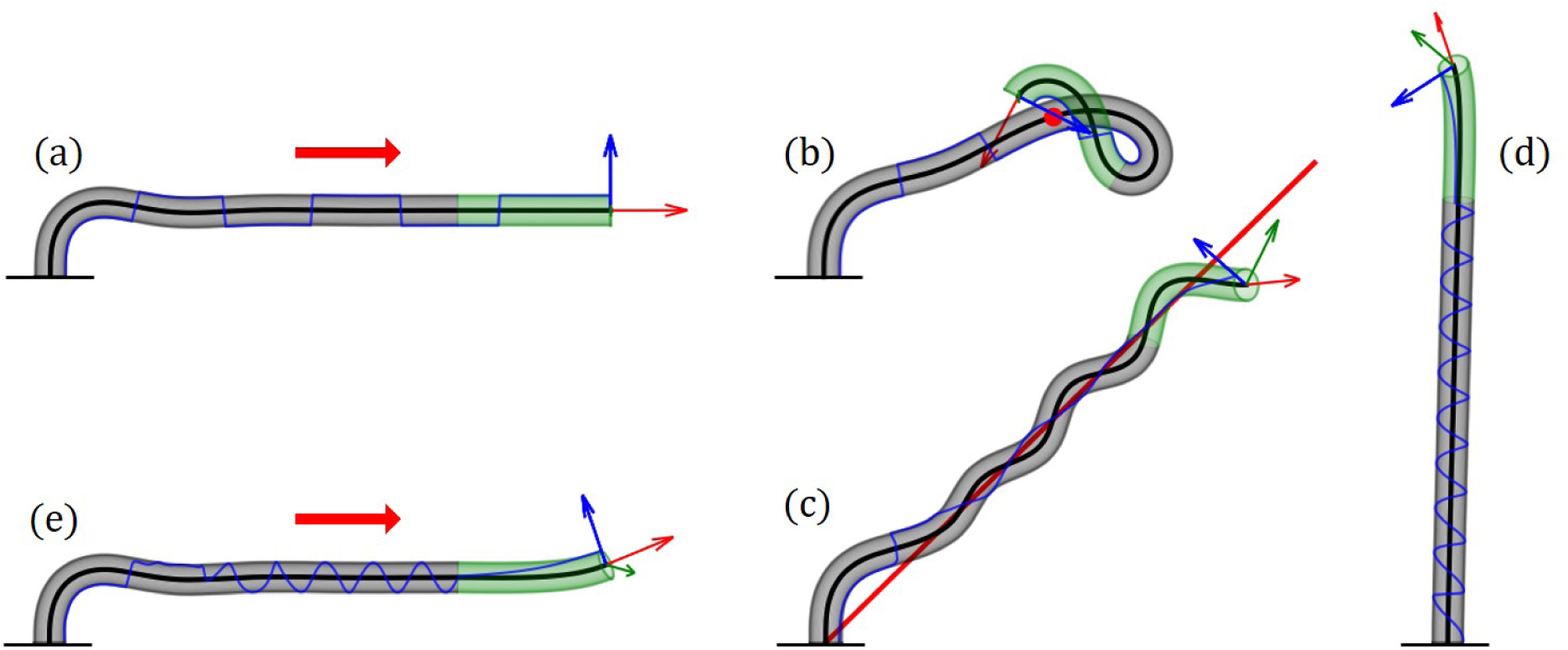
Examples of numerical simulations for various scenarios. Here we showcase snapshots of simulations for various cases. The subapical active growth zone is in green, while no growth occurs below that in grey. The arrows on the apex are the apical tangent direction 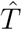 (red), normal direction 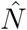 (blue) and the bi-normal direction 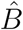 (green). The blue line marks the history of the direction of 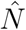 along the organ. The details of the simulations are brought in Sec. 3. We note that elasticity is not implemented here, and therefore the organ grows through itself. (a) Infinitely distant constant stimulus (red arrow). The organ reaches a steady state growing in the direction of the stimulus. 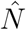 switches directions due to damped oscillations in the solution (Video 1 in SM). (b) Point stimulus (red dot). Illustrates the different dynamics between a distant vs. nearby stimulus (Video 2 in SM). (c) General geometry: twining around a line stimulus (red line). Any geometry for the source stimulus can be implemented. Here we chose a line geometry, which together with a signal in the direction of the line (to prevent self intersections) yields dynamics similar to twining of a climbing plant (Video 3 in SM). (d) Circumnutations. We implement the growth response to an internal cue rather than external cues, yielding inherent periodic movements of plants called circumnutations, generally associated with search processes. The periodic trajectory of 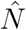 visualizes the rotational movement of the growing tip (Video 4 in SM). (e) Superposition of internal and external stimuli. We combine circumnutations together with an infinitely distant external stimulus (Video 5 in SM).

### Point stimulus

We consider the case of a stimulus whose source is a point located at 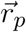 (Bastien et al., 2019), such as a nearby localized light or water source. In this case the stimulus leads to a radial vector field centered at the point, i.e. 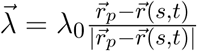:

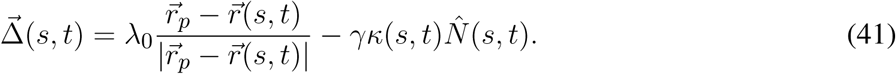

Here again *λ*_0_ is constant in space, however this can be generalized to depend on space, for example in the case of a diffusive chemical where 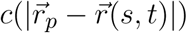. A snapshot of the dynamics is shown in Fig. 6b, while the full dynamics can be found in Video 2 in the SM.

### General stimulus geometry: twining around a line stimulus

We can generalize the point stimulus to any geometrical form. Here we show an example of a stimulus in the form of an attracting straight line. Let us assume the line is parallel to an arbitrary direction 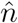, whose base position in the x-y plane is 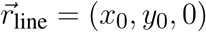. The minimal distance between a point on the organ 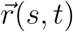 and the line is:

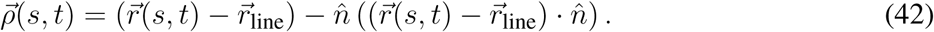

The response vector will then be 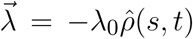. As an example of multiple stimuli, we also add a directional stimulus parallel to the line (i.e. gravity or light), 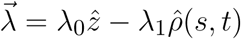, leading to the following differential growth vector :

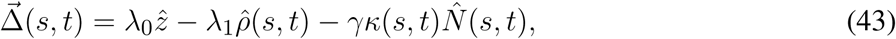

where 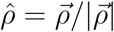. The resulting dynamics are reminiscent of the twining motion typical of climbing plants, as shown in Fig. 6c, and Video 3 in the SM. We note this twining movement is not based on touch, meaning that the organ does not hold the support. Furthermore no elasticity is involved at this stage, as further discussed in the Discussion section.

### Internal processes: circumnutations

Circumnutations are circular periodic movements of the tips of plant organs, generally associated with search processes, for example climbing plants searching for a support, or roots searching for nutrients. Unlike tropisms, these are inherent movements due to internal drivers, not external stimuli, and can be described as 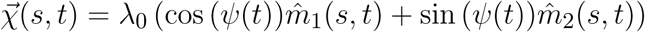 where *λ*_0_ is the intensity of the bending, *ψ*(*t*) is a general function describing the direction of growth at time *t*, and we described the direction of growth using the natural frame. Here we chose a circular form, however more elaborate forms can be used Bastien and Meroz (2016). Following Bastien and Meroz (2016), we substitute 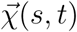 in the differential growth vector:

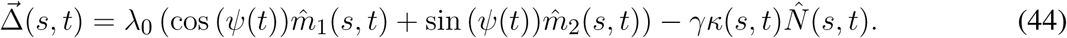

In our simulations, we took *ψ*(*t*) = *ωt* with *ω* = 0.2*/dt*. A snapshot is found in Fig. 6, and the full dynamics can be found in Video 4 in the SM. The trajectory of 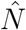 clearly illustrates the circular movement of the tip over time.

### Superposition of internal and external stimuli

As already suggested in the example of the line stimulus, where a directional stimulus was added, we can consider multiple types of stimuli by assuming they are additive. We bring here another example based on plant behavior, where we consider an organ responding to a distant external signal, while also exhibiting internally driven circumnutations. In this case we simply add to Eq. 44 the term for the distant stimulus in direction 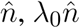, yielding:

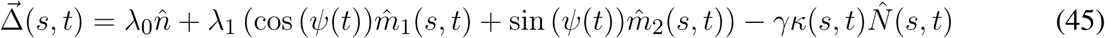

A snapshot of the resulting dynamics is shown in Fig. 6e, and the full dynamics are shown in Video 5 in the SM.

## 5 EXAMPLE OF AN OPTIMAL CONTROL APPROACH

In this last section we take a step back, and consider a simple example illustrating the possible use of control theory in order to recover tropic dynamics - in a way which may be relevant for robotics use. In what follows we no longer use the Frenet Serret formalism developed in the paper, relaxing the assumption of a constant arc-length parametrization. Instead we consider the general case where the curve of the organ is parametrized using the Lagrangian coordinate *S*_0_, as described in Section 2.2, without further reparametrizing the curve as it evolves over time. This general case may be pertinent to some robotics systems. We consider an organ with apical sensing, a fixed length L (neglecting an explicit account for growth as discussed before, and dynamics are restricted to 2D, similar to the case of apical sensing discussed in (Bastien et al., 2013). The aim is to find a controlled evolution equation of the tangent 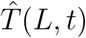 at the tip, where 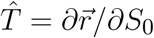. Let 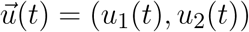 be a control to orient the tangent at the tip 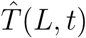. The sensing occuring at the tip influences the dynamics at any other point on the organ, and therefore 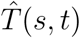 will satisify the following Cauchy problem for any s:

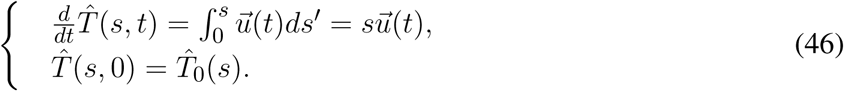

We further limit the family of possible control strategies to those for which:

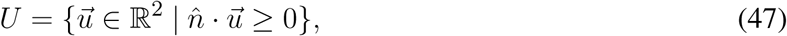

where 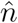 is the direction of the stimulus, since 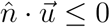 leads to undesired curling dynamics. From these strategies, we wish to choose those which are optimal in some sense. We therefore require that the optimal strategy minimizes some cost function which may manifest some physical element of the robot. Here we choose the following:

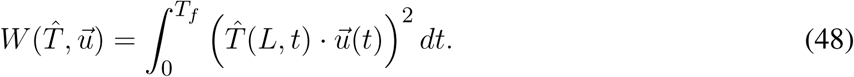

In this case the cost function has a geometric meaning: when the dot product goes to zero, together with Eq. 46 we have 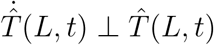, i.e. 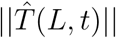 is constant, thus recovering the assumption at the basis of the Frenet-Serret formalism, of identical parametrization of the arc length over time. This gives a family of optimal controls:

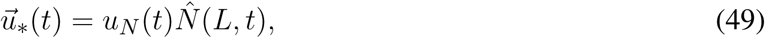

where 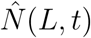 is the normal direction of the apex, and 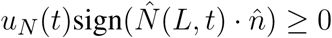 to satisfy Eq.47. These solutions ensures the tip approaches the stimulus in a strictly decreasing manner. Indeed, if the initial tangent 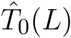 is not parallel to 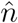, then

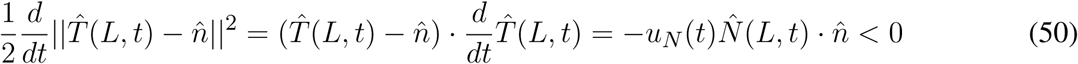

for all *t* such that 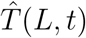 is not parallel to 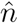. Then, 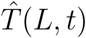 remains constantly parallel to 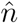. In particular, such a computation implies that the tangent 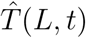 does not oscillate around the stimulus direction 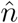. We focus our attention to a member of the control family described in Eq. 49:

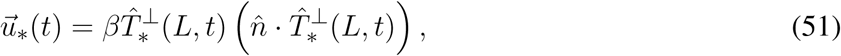

for all *t* ∈ [0, *T*_*f*_] and *β* ≥ 0, where 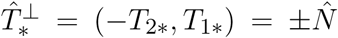 is just the perpendicular vector to 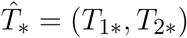. In Appendix C in the SM we show the details of the calculation based on the Pontryagin’s maximum principle (Aronna et al., 2017), showing that this indeed meets the requirements of the optimal control problem described in Eqs. 46, 47, 48. Substituting the specific solution described in Eq. 51 in the dynamics of Eq. 46, while also writing the tangent vector in terms of the angle *θ*(*s, t*) between the 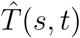 and the stimulus direction 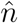, i.e. 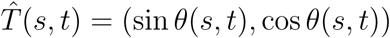, yields the following dynamical equation:

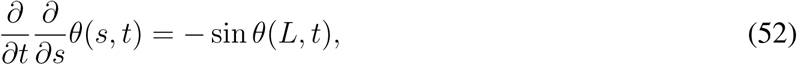

which is identical to the dynamics described in (Bastien et al., 2013), in the case of apical sensing, where proprioception is not required for stability.

## 6 DISCUSSION AND CONCLUSION

In this work we presented a general and rigorous mathematical framework of a rod-like growing organ, whose dynamics are driven by a differential growth vector. We construct the differential growth vector by taking into account both internal and external cues, as well as posture control, as schematically illustrated in Fig. 5. The model adopts the 3D Frenet-Serret formalism, which is a natural choice to describe curves, and useful for robotics control purposes.

In recent years there has been an advancement in the mathematical description of plant growth-driven movements. AC and ACE models (Bastien et al., 2013, 2014, 2015) describe 2D tropic dynamics in the case of an infinitely distant stimulus (e.g. gravitropism and phototropism), addressing the influence of growth as the driver of movement, and identifying the requirement of proprioception for stable dynamics. Bastien et al. (2019) generalizes this model in the case of a nearby point stimulus (generalizable to other geometries), relevant for chemical stimuli, or a nearby point light source. Bressan et al. (2017) developed a model for gravitropism, considering accretive growth rather than elongation, while achieving stable dynamics by controlling the growth-zone and sensitivity. Another model focuses on circumnutations in 3D (Bastien and Meroz, 2016), however disregards responses to external stimuli. A careful comparison of the models finds that our model is general, consolidating these different aspects in 3D for the first time: growth-driven responses to both external and internal cues, allowing stimuli with different physical and geometrical characteristics, while maintaining posture control through proprioception.

We ran numerical simulations of a number of key example cases. In the case of the response to external stimuli, we consider a distant stimulus (such as sunlight and gravity), a point stimulus (such as a point light source), and a rod stimulus which emulates *twining* of a climbing plant around a support. We also simulate circumnutations, the response to an internal oscillatory cue, associated with search processes. Lastly we also demonstrate the superposition of both the response to an external stimulus together with circumnutations. These examples showcase the broad spectrum of cases this framework can describe, and represent interactions with the environment which are at the basis of autonomous performance of next generation self-growing robots in unstructured scenarios, including movement in uncertain terrains involving obstacles and voids. The model presented here therefore sets the basis for a control system for robots with a changing and unpredictable morphology.

Though the framework we develop here successfully describes the various growth-driven movements of plants, it disregards parameters pertinent to robotic structures, such as energy, friction, weight etc. We bring a simple example illustrating the possible use of optimal control theory in order to recover tropic dynamics, in a way which may be relevant for robotics use. Optimal control theory optimizes processes where some cost function is minimized, and is therefore useful in engineering problems. The example *per se* does not necessarily present a practical cost function, however it suggests that future work may include optimizing the current model for tropic movements so as to minimize a cost function associated with a robotic parameter.

This general framework allows a better understanding of plant tropisms and plant behavior, which has only recently become the focus of research. Basic behavioral processes in animals are generally studied through the motor responses to controlled stimuli, and a solid understanding of plant movements (both in response to internal and external cues) paves the way to designing controlled behavioral experiments. For example, simulations of incorporating both circumnutations and tropisms will allow to quantitatively investigate the role of circumnutations in the the successful search after nutrients or light.

Furthermore, careful attention has been put in relating environmental stimuli to differential growth, discussing stimuli with different physical characteristics, categorized by their mathematical description, such as scalar fields (concentration of water or nutrients) and vector fields (light and gravity). This analysis is important from the robotics standpoint, and it also advances our understanding in plant tropisms, allowing a better characterization of plant biosensors in tropisms which are less understood, such as hydrotropism and chemotropism. We note that though this framework is inspired by plant responses, it is not based on biological details, and is therefore amenable to any rod-like organisms which respond to signals via growth, such as neurons and fungi.

Lastly we note that this framework does not currently include mechanics or elasticity, disregarding any elastic responses of the organ to physical forces. However this can be naturally implemented in the Frenet-Serret frame of reference (Goriely, 2017; Chelakkot and Mahadevan, 2017; Agostinelli et al., 2020), which we plan in future work.

## AUTHOR CONTRIBUTIONS

AP and YM developed the mathematical framework, AP carried out the numerical analysis and simulations, and prepared all figures and videos. FT, MP and PM developed the optimal control calculation, and the comparison with the model by Bressan et al. AP and YM drafted the manuscript. All authors contributed to the authoring of the final manuscript and contributed to the research design.

## ACKNOWLEDGMENTS

We thank Renaud Bastien for helpful conversations.

## SUPPLEMENTAL DATA

The supplementary material includes one pdf file, and 5 videos.

**SM.pdf** includes three Appendices:

Appendix A: Derivation of growth dynamics both in the Frenet-Serret frame and the natural frame, relevant to Sections 2.2 and 3.

Appendix B: shows that our 3D simulations converge to the known analytical solution in 2D with an exponential growth profile.

Appendix C: Pontryagin’s Maximum Principle, part of the calculation in the optimal control approach in Section 5.

The videos show simulations for the different cases presented in Section 4, and Fig. 6 shows snapshots of the simulations. The arrows represent the Frenet-Serret Frame (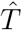 in red, 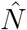 in blue, and 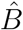 in green). The graphs show the values of the curvature *κ*(*s, t*) and torsion *ϕ*(*s, t*) as a function of time:

**Video1.mp4** Infinitely distant constant stimulus.

**Video2.mp4** Point stimulus.

**Video3.mp4** General stimulus geometry: twining around a line stimulus.

**Video4.mp4** Internal processes: circumnutations.

**Video5.mp4** Superposition of internal and external stimuli.

## DATA AVAILABILITY STATEMENT

The numerical code for simulations presented in this work are accessible at: https://github.com/poratamir/3D-growth-dynamics

## REFERENCES

Agostinelli, D., Lucantonio, A., Noselli, G., and DeSimone, A. (2020). Nutations in growing plant shoots: the role of elastic deformations due to gravity loading. Journal of mechanics and physics of solids 136. doi: 10.1016/j.jmps.2019.103702

Aronna, M. S., Tonon, D., Boccia, A., Campos, C. M., Mazzola, M., Van Nguyen, L., et al. (2017). Optimality conditions (in pontryagin form). In Optimal Control: Novel Directions and Applications (Springer). 1–125

Bastien, R., Bohr, T., Moulia, B., and Douady, S. (2013). Unifying model of shoot gravitropism reveals proprioception as a central feature of posture control in plants. Proceedings of the National Academy of Sciences of the United States of America 110, 755–760

Bastien, R., Douady, S., and Moulia, B. (2014). A unifying modeling of plant shoot gravitropism with an explicit account of the effects of growth. Frontiers in Plant Science 5

Bastien, R., Douady, S., and Moulia, B. (2015). A Unified Model of Shoot Tropism in Plants: Photo-, Gravi- and Propio-ception. PLoS Comput Biol 11, e1004037

Bastien, R. and Meroz, Y. (2016). The Kinematics of Plant Nutation Reveals a Simple Relation between Curvature and the Orientation of Differential Growth. PLoS Comput Biol 12, e1005238

Bastien, R., Porat, A., and Meroz, Y. (2019). Towards a framework for collective behavior in growth-driven systems, based on plant-inspired allotropic pairwise interactions. Bioinspir. Biomim. 14

Bishop, R. L. (1975). There is more than one way to frame a curve. The American Mathematical Monthly 82, 246–251. doi: 10.1080/00029890.1975.11993807

Bressan, A., Palladino, M., and Shen, W. (2017). Growth models for tree stems and vines. Journal of Differential Equations 263, 2280 – 2316. doi: 10.1016/j.jde.2017.03.047

Chauvet, H., Pouliquen, O., Forterre, Y., Legué, V., and Moulia, B. (2016). Inclination not force is sensed by plants during shoot gravitropism. Scientific Reports 6, 35431

Chelakkot, R. and Mahadevan, L. (2017). On the growth and form of shoots. Journal of The Royal Society Interface 14, 20170001

Darwin, C. (1880). The Power of Movement in Plants. (London: John Murray Publishers)

Dottore, E. D., Mondini, A., Sadeghi, A., Mattoli, V., and Mazzolai, B. (2016). Circumnutations as a penetration strategy in a plant-root-inspired robot. 2016 IEEE International Conference on Robotics and Automation (ICRA), 4722–4728 doi: 10.1109/icra.2016.7487673

Forterre, Y. (2013). Slow, fast and furious: understanding the physics of plant movements. Journal of experimental botany 64, 4745–60. doi: 10.1093/jxb/ert230

Gazzola, D. L. M. A. M and L, M. (2018). Forward and inverse problems in the mechanics of soft filaments. Royal Society Open Science 5

Gilroy, S. and Masson, P. (2007). Plant Tropisms (Wiley-Blackwell)

Goriely, A. (2017). The Mathematics and Mechanics of Biological Growth (Springer)

Hamant, O. and Moulia, B. (2016). How do plants read their own shapes? New Phytologist 212, 333–337

Hawkes, E. W., Blumenschein, L. H., Greer, J. D., and Okamura, A. M. (2017). A soft robot that navigates its environment through growth. Science Robotics 2. doi: 10.1126/scirobotics.aan3028

Hohm, T., Preuten, T., and Fankhauser, C. (2013). Phototropism: Translating light into directional growth. American Journal of Botany 100, 47–59

Holland, J., Roberts, D., and Liscum, E. (2009). Understanding phototropism: from darwin to today. Journal of Experimental Botany 60, 1969–1978

Knieb, E., Salomon, M., and Rdiger, W. (2004). Tissue-specific and subcellular localization of phototropin determined by immuno-blotting. Planta 218, 843–851

Langer, J. and Singer, D. A. (1996). Lagrangian aspects of the kirchhoff elastic rod. SIAM Review 38, 605–618. doi: 10.1137/S0036144593253290

Laschi, C. and Mazzolai, B. (2016). Lessons from Animals and Plants: The Symbiosis of Morphological Computation and Soft Robotics. IEEE Robotics & Automation Magazine 23, 107–114

Mazzolai, B. (2016). Plant-Inspired Growing Robots (Springer International Publishing), Soft Robotics: Trends, Applications and Challenges. 57 – 63

Mazzolai, B., Mattoli, V., and Beccai, L. (2016). Soft Plant Robotic Solutions: Biological Inspiration and Technological Challenges (Springer International Publishing), vol. 23. 8 edn., 687 – 707

Mehling, J. S., Diftler, M. A., Chu, M., and Valvo, M. (2006). A Minimally Invasive Tendril Robot for In-Space Inspection. The First IEEE/RAS-EMBS International Conference on Biomedical Robotics and Biomechatronics, 2006. BioRob 2006., 690–695 doi: 10.1109/biorob.2006.1639170

Morita, M. T. and Tasaka, M. (2004). Gravity sensing and signaling. Current Opinion in Plant Biology 7, 712–718

Norwich, K. and Wong, W. (1997). Unification of psychophysical phenomena: The complete form of fechner’s law. Perception and Psychophysics doi: 10.3758/BF03205509

Rivière, M., Derr, J., and Douady, S. (2017). Motions of leaves and stems, from growth to potential use. Physical biology 14, 051001

Sadeghi, A., Mondini, A., and Mazzolai, B. (2017). Toward Self-Growing Soft Robots Inspired by Plant Roots and Based on Additive Manufacturing Technologies. Soft Robotics 4, 211–223

Sadeghi, A., Tonazzini, A., Popova, L., and Mazzolai, B. (2014). A Novel Growing Device Inspired by Plant Root Soil Penetration Behaviors. PLOS ONE 9, e90139

Sakamoto, K. and Briggs, W. (2002). Cellular and subcellular localization of phototropin 1. The Plant Cell Online 14, 1723–1735

Silk, W. K. (1989). Growth Rate Patterns which Maintain a Helical Tissue Tube. J. theor. Biol. 138, 311–327

Stevens, S. (1957). On the psychophysical law. Psychological Review doi: 10.1037/h0046162

Su, S.-H., Gibbs, N. M., Jancewicz, A. L., and Masson, P. H. (2017). Molecular Mechanisms of Root Gravitropism. Current Biology 27, R964–R972

Wan, Y., Eisinger, W., Ehrhardt, D., Kubitscheck, U., and Baluska, F. e. a. (2008). The subcellular localization and blue-light-induced movement of phototropin 1-gfp in etiolated seedlings of arabidopsis thaliana. Molecular Plant 1, 103–117

