## Supplementary Material for "A general 3D model for growth dynamics of sensory-growth systems: from plants to robotics"

#### APPENDIX A: DERIVATION OF THE GROWTH DYNAMICS

We would like to model the active reorientation of organs as a result of local differential growth. Here, we define the differential growth of our cylindrical organs as a non-uniform growth rate within a cross section of an organ, that keeps the cross-section of the organ circular with a constant radius. This an be achieved by letting the organ grow only parallel to the centerline, and by expressing the growth rates in every cross-section using only properties of the centerline. We start by focusing our attention on an infinitesimal part of centerline, defined as:

$$ds(S_0,t) = s(S_0 + dS_0, t) - s(S_0, t) \quad . \tag{S1}$$

Using the relation:

$$\frac{ds}{dt} = v(s,t) = \int_0^s \dot{E}(u,t)du \quad , \tag{S2}$$

the time derivative of the segment can be written as:

$$\frac{d}{dt}ds(S_0,t) = \frac{d}{dt}\left(s(S_0 + dS_0, t) - s(S_0,t)\right) = \int_{s(S_0,t)}^{s(S_0+dS_0,t)} \dot{E}(u,t)du \approx \tag{S3}$$
$$\approx \dot{E}(s(S_0,t),t)\left(s(S_0 + dS_0, t) - s(S_0,t)\right) = \dot{E}(s(S_0,t),t)ds(S_0,t) \quad .$$

Omitting the dependence of variables on $(S_0, t)$ or $(s, t)$, Eq. S3 gives an expression of the local growth rate of the centerline:

$$\frac{1}{ds}\frac{d}{dt}ds = \dot{E} \tag{S4}$$

We now turn to calculate the growth rates of lines parallel to the centerline on the surface of the organ. For this, a local coordinate frame is required. We now show how to reach the dynamic equations using both the Frenet-Serret frame and the natural frame.

##### Using the Frenet-Serret Frame

We begin by deriving the dynamics of the growing rod using the Frenet-Serret formalism. This can be done when the curvature of the centerline doesn't nullify, and the Frenet-Serret coordinate system is well defined on the centerline. We then take inspiration from (Silk, 1989) and describe lengths of lines on the surface of the organ which are parallel to the centerline. To do so, we stretch a vector to a point on the perimeter of the cross section using cylindrical coordinates $(\rho, \phi)$:

$$\vec{r} = \vec{r}_0 + R\left(\cos(\phi)\hat{N} - \sin(\phi)\hat{B}\right) \equiv \vec{r}_0 + R\hat{\rho} \tag{S5}$$

Where $\vec{r}_0$ is the centerline, $R$ is the radius of the organ, and $\phi$ is the angle of the point with respect to the normal direction $\hat{N}$. In the last equality we defined the unit vector $\hat{\rho}$ for later use. We note that all of the variables appearing in Eq. S5 have an implicit dependence on the arc length and time, $(s, t)$. The lengths of lines parallel to the centerline can be found using the first fundamental form:

$$\ell(\hat{\rho}) = \left|\frac{\partial \vec{r}}{\partial s}\right| ds \tag{S6}$$

Using Eq. S5, the definition of $\hat{T}$ and the Frenet-Serret equations, we find:

$$\begin{aligned}\frac{\partial \vec{r}}{\partial s} &= \frac{\partial \vec{r}_0}{\partial s} + R\frac{\partial}{\partial s}\left(\cos(\phi)\hat{N} - \sin(\phi)\hat{B}\right) = \\ &= \hat{T} + R\left(\cos(\phi)(-\kappa\hat{T} + \tau\hat{B}) - \sin(\phi)(-\tau\hat{N})\right) \\ &\qquad - R\left(\sin(\phi)\frac{\partial\phi}{\partial s}\hat{N} + \cos(\phi)\frac{\partial\phi}{\partial s}\hat{B}\right) = \\ &= \hat{T}(1 - R\kappa\cos(\phi))\end{aligned} \tag{S7}$$

Where in the last equality we used the relation:

$$\frac{\partial\phi}{\partial s} = \tau \tag{S8}$$

Inserting Eq. S7 into Eq. S6 gives:

$$\ell(\hat{\rho}) = (1 - R\kappa\cos(\phi))ds = (1 - R\kappa\hat{N}\cdot\hat{\rho})ds \tag{S9}$$

Where we assumed the low curvature limit: $R\kappa \ll 1$. Now, similar to Eq. S4, the growth rate of the lines on the surface of the organ which are parallel to the centerline can be obtained by:

$$\dot{\epsilon}(\hat{\rho}) = \frac{1}{\ell(\hat{\rho})}\frac{d\ell(\hat{\rho})}{dt} \tag{S10}$$

We now write this time derivative explicitly. We take the time derivative of $\ell(\hat{\rho})$ using Eq. S9, use Eq. S4 for the time derivative of $ds$, and the material derivative for the Lagrangian quantities $\kappa, \hat{N}$. All together, this gives:

$$\dot{\epsilon}_R(\hat{\rho}) = \dot{E} - \frac{1}{1 - R\kappa\hat{N}\cdot\hat{\rho}}\frac{D(R\kappa\hat{N}\cdot\hat{\rho})}{Dt} \approx \dot{E} - R\frac{D(\kappa\hat{N}\cdot\hat{\rho})}{Dt}, \tag{S11}$$

where in the last equality we took once more the low curvature limit: $R\kappa \ll 1$. We note that the growth rates in Eq. S11 can describe completely the growth in the entire volume of the cylinder, by changing $R$ to a smaller radius $\rho$ such that $0 \leq \rho \leq R$. The growth everywhere is thus governed by the properties of the centerline, and is tailored to maintain a constant circular cross section. We now define a measure of the differential growth within the cross section. We note the differential growth towards the $\hat{\rho}$ direction on the $(\hat{N}, \hat{B})$ plane as the difference in growth rate on either side of $\hat{\rho}$, normalized by their sum:

$$\Delta(\hat{\rho}) \equiv \frac{\dot{\epsilon}_R(-\hat{\rho}) - \dot{\epsilon}_R(\hat{\rho})}{\dot{\epsilon}_R(-\hat{\rho}) + \dot{\epsilon}_R(\hat{\rho})} = \frac{R}{\dot{E}}\frac{D(\kappa\hat{N}\cdot\hat{\rho})}{Dt} \tag{S12}$$

Using this definition, if $\Delta(\hat{\rho}) > 0$, the organ grows faster on the $-\hat{\rho}$ direction than the $\hat{\rho}$ direction, which will lead to bending to the $\hat{\rho}$ direction. We now define a vector of differential growth in the local cross section plane, which points to the maximal growth direction:

$$\vec{\Delta} = \Delta(\hat{\rho} = \hat{N})\hat{N} + \Delta(\hat{\rho} = \hat{B})\hat{B} \tag{S13}$$

To receive the final form of the dynamics equations, we write Eq. S12 using the angle $\phi$ by noticing that $\hat{N} \cdot \hat{\rho} = \cos(\phi)$. This gives:

$$\Delta_\phi(\phi) = \frac{R}{\dot{E}} \frac{D(\kappa \cos(\phi))}{Dt} = \frac{R}{\dot{E}} \left( \frac{D\kappa}{Dt} \cos(\phi) - \kappa \sin(\phi) \frac{D\phi}{Dt} \right) \tag{S14}$$

Then, using Eq. S13 and Eq. S14, we note that:

$$\vec{\Delta} \cdot \hat{N} = \Delta_\phi(\phi = 0) = \frac{R}{\dot{E}} \frac{D\kappa}{Dt}, \tag{S15}$$

$$\vec{\Delta} \cdot \hat{B} = \Delta_\phi \left( \phi = -\frac{\pi}{2} \right) = \frac{R}{\dot{E}} \kappa \frac{D\phi}{Dt}, \tag{S16}$$

which are the equations as appeared in the main text:

$$\boxed{\begin{aligned} \frac{D}{Dt}\kappa &= \frac{\dot{E}}{R} \vec{\Delta} \cdot \hat{N} \\ \kappa \frac{D}{Dt}\phi &= \frac{\dot{E}}{R} \vec{\Delta} \cdot \hat{B} \end{aligned}} \tag{S17}$$

Therefore, if we give an expression for the differential growth vector $\vec{\Delta}(s, t)$ and an initial configuration, the dynamics of the entire rod can be integrated completely.

#### Using the natural frame

We now place the "natural frame" $(\hat{m}_1, \hat{m}_2, \hat{T})$ on the centerline. This frame is well defined even when the curvature nullifies. To continue, we describe again the lengths of lines which are parallel to the centerline. To do so, we stretch a vector to a point on the cross section:

$$\vec{r} = \vec{r}_0 + x_1 \hat{m}_1 + x_2 \hat{m}_2 \tag{S18}$$

Where $\vec{r}_0$ is the centerline and $(x_1, x_2)$ are the Cartesian coordinates in the cross section plane. We note that $x_1^2 + x_2^2 \leq R^2$, since the cross section is circular. The lengths of the lines which are parallel to the centerline can be found using the first fundamental form:

$$\ell(x_1, x_2) = \left| \frac{\partial \vec{r}}{\partial s} \right| ds \tag{S19}$$

Plugging the expression for $\vec{r}$ from Eq. S18 into Eq. S19 and using the natural frame's relations for the derivatives (as appears in the main text), we obtain that:

$$\ell(x_1, x_2) = \left| \frac{\partial \vec{r}}{\partial s} \right| ds = \left| \hat{T} - x_1 \kappa_1 \hat{T} - x_2 \kappa_2 \hat{T} \right| ds = (1 - x_1 \kappa_1 - x_2 \kappa_2) ds \tag{S20}$$

Where in the last equality we assume: $R\kappa = R\sqrt{\kappa_1^2 + \kappa_2^2} \ll 1$, which gives $\kappa_1 x_1 \ll 1$ and $\kappa_2 x_2 \ll 1$. We continue by calculating the growth rates of the lines parallel to the centerline using:

$$\dot{\epsilon}(x_1, x_2) = \frac{1}{\ell(x_1, x_2)} \frac{d}{dt} \ell(x_1, x_2), \tag{S21}$$

and assuming that the curvatures of the centerline are time dependent. Since they are Eularian quantities (with respect to the growth flow), a material derivative is needed on the curvatures. Deriving Eq. S20 with respect to time, and using Eq. S4, we get:

$$\frac{d}{dt}\ell(x_1, x_2) = (1 - x_1\kappa_1 - x_2\kappa_2)\dot{E}ds - (x_1 \frac{D\kappa_1}{Dt} + x_2 \frac{D\kappa_2}{Dt})ds \tag{S22}$$

Plugging Eq. S22 into Eq. S21 gives:

$$\dot{\epsilon}(x_1, x_2) = \dot{E} - \frac{x_1 \frac{D\kappa_1}{Dt} + x_2 \frac{D\kappa_2}{Dt}}{1 - x_1\kappa_1 - x_2\kappa_2} \approx \dot{E} - x_1 \frac{D\kappa_1}{Dt} - x_2 \frac{D\kappa_2}{Dt} \tag{S23}$$

Where in the last equality we assumed once again that $R\kappa = R\sqrt{\kappa_1^2 + \kappa_2^2} \ll 1$.
The growth rates on the surface of the cylindrical organ can be written using cylindrical coordinates. If we mark the radial direction in the $(\hat{m}_1, \hat{m}_2)$ plane as $\hat{\rho}$, we can write:

$$x_1 = R\hat{m}_1 \cdot \hat{\rho} \tag{S24}$$

$$x_2 = R\hat{m}_2 \cdot \hat{\rho} \tag{S25}$$

$$\dot{\epsilon}_R(\hat{\rho}) = \dot{E} - R\hat{\rho} \cdot \left( \frac{D\kappa_1}{Dt}\hat{m}_1 + \frac{D\kappa_2}{Dt}\hat{m}_2 \right) \tag{S26}$$

We now define a measure of the differential growth within the cross section. We note the differential growth towards the $\hat{\rho}$ direction on the $(\hat{m}_1, \hat{m}_2)$ plane as the difference in growth rate on either side of $\hat{\rho}$ at the surface of the organ, normalized by their sum:

$$\Delta(\hat{\rho}) \equiv \frac{\dot{\epsilon}_R(-\hat{\rho}) - \dot{\epsilon}_R(\hat{\rho})}{\dot{\epsilon}_R(-\hat{\rho}) + \dot{\epsilon}_R(\hat{\rho})} = \frac{R}{\dot{E}}\hat{\rho} \cdot \left( \frac{D\kappa_1}{Dt}\hat{m}_1 + \frac{D\kappa_2}{Dt}\hat{m}_2 \right) \tag{S27}$$

Using this definition, if $\Delta(\hat{\rho}) > 0$, the organ grows faster on the $-\hat{\rho}$ direction than the $\hat{\rho}$ direction, which will lead to bending to the $\hat{\rho}$ direction. We now define a vector of differential growth in the local cross section plane, which will point to the maximal growth direction:

$$\vec{\Delta} = \Delta(\hat{m}_1)\hat{m}_1 + \Delta(\hat{m}_2)\hat{m}_2 = \frac{R}{\dot{E}} \left( \frac{D\kappa_1}{Dt}\hat{m}_1 + \frac{D\kappa_2}{Dt}\hat{m}_2 \right) \tag{S28}$$

We can thus use the differential growth vector in order to describe the dynamics of local curvature:

$$\boxed{\begin{aligned}\frac{D\kappa_1}{Dt} &= \frac{\dot{E}}{R}\vec{\Delta}\cdot\hat{m}_1 \\ \frac{D\kappa_2}{Dt} &= \frac{\dot{E}}{R}\vec{\Delta}\cdot\hat{m}_2\end{aligned}} \tag{S29}$$

Therefore, if we give an expression for the differential growth vector $\vec{\Delta}$ and an initial configuration, the dynamics of the entire rod can be integrated completely.
Assuming $\kappa \neq 0$, we can translate the resulting dynmamics of Eq. S29 to the Frenet-Serret frame, and express them using $\kappa, \phi, \hat{N}$ and $\hat{B}$. Given the relations:

$$\kappa = \sqrt{\kappa_1^2 + \kappa_2^2} \tag{S30}$$

$$\phi = \arctan\left(\frac{\kappa_2}{\kappa_1}\right) \tag{S31}$$

$$\hat{N} = \cos(\phi)\hat{m}_1 + \sin(\phi)\hat{m}_2 \tag{S32}$$

$$\hat{B} = -\sin(\phi)\hat{m}_1 + \cos(\phi)\hat{m}_2 \tag{S33}$$

$$\tag{S34}$$

We find that:

$$\frac{D\kappa}{Dt} = \frac{D}{Dt}\sqrt{\kappa_1^2 + \kappa_2^2} = \tag{S35}$$

$$= \frac{\kappa_1}{\sqrt{\kappa_1^2+\kappa_2^2}}\frac{D\kappa_1}{Dt} + \frac{\kappa_2}{\sqrt{\kappa_1^2+\kappa_2^2}}\frac{D\kappa_2}{Dt} = \tag{S36}$$

$$= \frac{\dot{E}}{R}\left(\cos(\phi)\vec{\Delta}\cdot\hat{m}_1 + \sin(\phi)\vec{\Delta}\cdot\hat{m}_2\right) = \tag{S37}$$

$$= \frac{\dot{E}}{R}\vec{\Delta}\cdot\hat{N} \tag{S38}$$

and:

$$\frac{D\phi}{Dt} = \frac{D}{Dt}\arctan\left(\frac{\kappa_2}{\kappa_1}\right) = \tag{S39}$$

$$= \frac{1}{1+\left(\frac{\kappa_2}{\kappa_1}\right)^2}\left(\frac{1}{\kappa_1}\frac{D\kappa_2}{Dt} - \frac{\kappa_2}{\kappa_1^2}\frac{D\kappa_1}{Dt}\right) = \tag{S40}$$

$$= \frac{\kappa_1}{\kappa_1^2+\kappa_2^2}\frac{D\kappa_2}{Dt} - \frac{\kappa_2}{\kappa_1^2+\kappa_2^2}\frac{D\kappa_1}{Dt} = \tag{S41}$$

$$= \frac{\dot{E}}{R\kappa}\left(\cos(\phi)\vec{\Delta}\cdot\hat{m}_2 - \sin(\phi)\vec{\Delta}\cdot\hat{m}_1\right) = \tag{S42}$$

$$= \frac{\dot{E}}{R\kappa}\vec{\Delta}\cdot\hat{B} \tag{S43}$$

And the two derivations agree.

### APPENDIX B: NUMERICAL ACCURACY VALIDATION

In *Bastien 2014 Front. Plant Sci*, a 2-d analysis of growing organs is presented. There, an organ that grows at every arc length is actively responding to gravity with a proprioception term. In constant stimuli, the shape of the organ is restricted to the plane defined by the initial orientation of the organ and the direction of gravity, as explained in the main text. In order to find the organ's dynamics, some simplifications are assumed:

- Constant growth rate all along the organ: $\dot{E}(s,t) = \text{const}$. This gives a simple growth velocity profile: $v(s,t) = s\dot{E}$, which is also known as an exponential growth profile.
- Writing the active growth equation using the local angle with respect to gravity. This angle can be calculated by:

$$\theta(s,t) = \int_0^s \kappa(s',t)ds' - \theta_g \tag{S44}$$

  Where $\theta_g$ is the direction of gravity. The dynamics of the organ can then be written as:

$$\frac{D}{Dt}\left(\frac{\partial\theta}{\partial s}\right) = \frac{\partial^2\theta}{\partial s\partial t} + v\frac{\partial^2\theta}{\partial s^2} = \frac{\dot{E}}{R}\left(-\lambda\sin(\theta) - \gamma\frac{\partial\theta}{\partial s}\right) \tag{S45}$$

- Taking the small angles limit: If the organ's initial direction with respect to gravity is small ($\theta \ll 1$) the resulting differential equation becomes linear and analytically solvable:

$$\frac{\partial^2\theta}{\partial s\partial t} + s\dot{E}\frac{\partial^2\theta}{\partial s^2} = \frac{\dot{E}}{R}\left(-\lambda\theta - \gamma\frac{\partial\theta}{\partial s}\right) \tag{S46}$$

Using the above assumptions, the steady state solution ($\frac{\partial^2\theta}{\partial s\partial t} = 0$) for an initially straight rod organ pointed to the angle $\theta_0$ is:

$$\theta(s) = \theta_0\Gamma\left(\frac{\gamma}{R}\right)\left(\frac{\lambda s}{R}\right)^{\frac{1}{2}-\frac{\gamma}{2R}} J_{\frac{\gamma}{R}-1}\left(2\sqrt{\frac{\lambda s}{R}}\right) \tag{S47}$$

The curvature is then:

$$\kappa(s) = \frac{\partial\theta}{\partial s} = -\frac{\lambda\theta_0}{R}\Gamma\left(\frac{\gamma}{R}\right)\left(\frac{\lambda s}{R}\right)^{-\frac{\gamma}{2R}} J_{\frac{\gamma}{R}}\left(2\sqrt{\frac{\lambda s}{R}}\right) \tag{S48}$$

We validate our numerical method using simulations of this analytically solvable problem. We take organs with an initial length $L_0 = 1$, radius $R = 0.1$, proprioceptive coefficient $\gamma = 0.1$ and gravitropic coefficient of $\lambda = 1$. The initial angle with respect to gravity is $\theta_0 = 5\pi/180$, and the discretization step size of the centerline is varied: $ds = 0.1, 0.05, 0.01, 0.005$. Since the organ grows exponentially, we simulate only the initial length of the organ, ignoring it's total length. We iterate the dynamics over 1500 growth timesteps, using $dt = 0.1$, while normalizing the growth rate for each simulation using the relation $\dot{E} = ds/(L_0 dt)$. The results of the different simulations are presented in Fig. S1. To quantify the error for each discretization step size, we used:

$$\text{ERR} = \int_0^{L_0} (\kappa_{\text{sim}}^2 - \kappa_{\text{theory}}^2)ds \tag{S49}$$

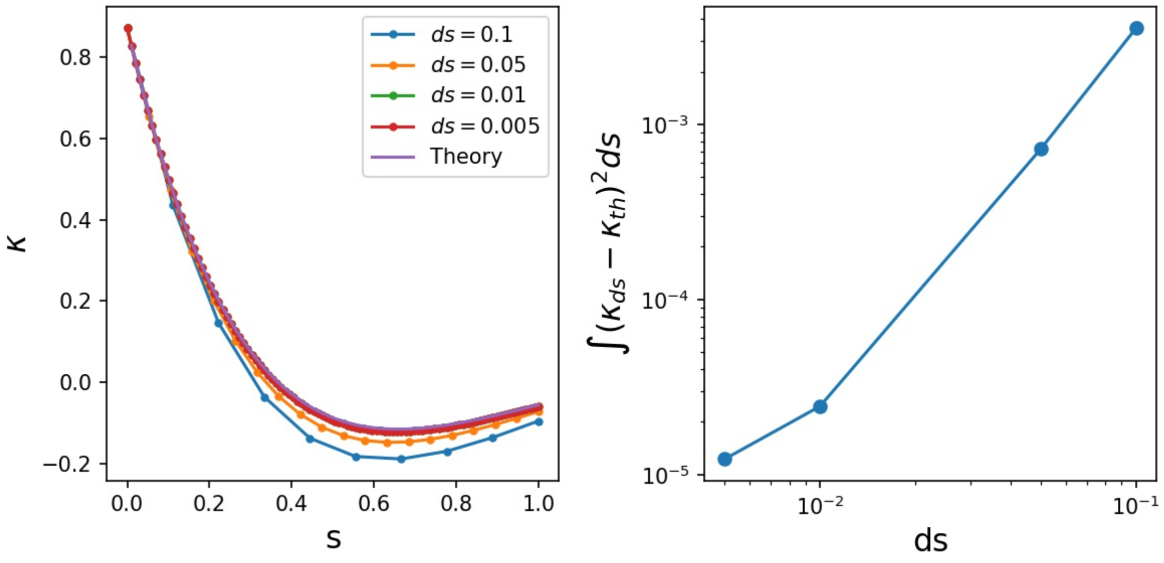


**Figure S1. Numerical accuracy validation:** Left: Results of simulations of the dynamics governed by Eq. S46. Each line shows the curvature as a function of arc-length for simulations with a different discretization step size $ds$. The simulation coverge towards the theoretical value from equation Eq. S48, which is plotted as a continuous line. Note that the dynamics are calculated only within the initial length of the organ. Right: Quantification of the error at the end of the simulations as a function of the discretization step size $ds$, as described by Eq. S49.

#### APPENDIX C: PONTRYAGIN'S MAXIMUM PRINCIPLE

Let $\vec{u}(t) = (u_1(t), u_2(t))$ be the control to orient the tangent at the tip $\hat{T}(L,t)$. The sensing occuring at the tip influences the dynamics of $\hat{T}(s,t)$ for all $s \in [0, L]$, accordingly to the following Cauchy problem:

$$\begin{cases} \frac{d}{dt}\hat{T}(s,t) = \int_0^s \vec{u}(t)ds' = s\vec{u}(t), \\ \hat{T}(s,0) = \hat{T}_0(s). \end{cases} \tag{S50}$$

To be optimal, the control $\vec{u}(t)$ has to satisfy all the conditions described in the main paper and summarised in the following optimal problem $(P)$:

$$(P) \begin{cases} \text{minimize}\, W(\hat{T}, \vec{u}) \\ \text{over } (\hat{T}, \vec{u})(\cdot) \in \mathrm{AC}([0, T_f]; \mathbb{R}^2) \times \mathcal{U}; \\ \frac{d}{dt}\hat{T}(L,t) = L\vec{u}(t) \text{ a.e. } t \in [0, T_f]; \\ \vec{u}(t) \in U \text{ a.e. } t \in [0, T_f]; \\ \hat{T}(L,0) = \hat{T}_0(L). \end{cases} \tag{S51}$$

$\mathrm{AC}([0, T_f]; \mathbb{R}^2)$ stands for absolutely continuous functions from $[0, T_f]$ to $\mathbb{R}^2$. $\mathcal{U}$ is the set of Lebesgue measurable functions defined on $[0, T_f]$ and taking values in $U = \{\vec{u} \in \mathbb{R}^2 \mid \hat{n} \cdot \vec{u} \geq 0\}$ and $\hat{n}$ is the direction of an approaching stimulus. To solve the problem $(P)$ we have to identify an optimal control $\vec{u}_*(t)$ (and hence the corresponding trajectory $\hat{T}_*(L,t)$) such that the couple $(\hat{T}_*, \vec{u}_*)(\cdot)$ minimizes the cost function

$$W(\hat{T},\vec{u}) = \int_0^{T_f} \left(\hat{T}(L,t)\cdot\vec{u}(t)\right)^2 dt. \tag{S52}$$

As a first step to solve $(P)$, one could use the so called *Pontryagin's maximum principle* (Aronna et al., 2017). It states necessary conditions the optimal control has to satisfy along the optimal state trajectory. To this aim, let us define the Hamiltonian $H$

$$H(\vec{T},\vec{\omega},\vec{p}) = L\,\vec{p}\cdot\vec{\omega} - \left(\vec{T}\cdot\vec{\omega}\right)^2, \tag{S53}$$

for any $(\vec{T},\vec{\omega},\vec{p}) \in \mathbb{R}^2 \times U \times \mathbb{R}^2$. The Pontryagin's maximum principle states that the state trajectory $T_*(t)$ and the control $\vec{u}_*(t)$ have to satisfy the following conditions to be optimal

$$\begin{cases} -\dot{\vec{p}}(t) = \partial_{\vec{T}} H(\hat{T}_*(L,t),\vec{u}_*(t),\vec{p}(t)) = -\vec{u}_*(t)\left(\hat{T}_*(L,t)\cdot\vec{u}_*(t)\right); \\ -\vec{p}(T_f) = 0; \\ H(\hat{T}_*(L,t),\vec{u}_*(t),\vec{p}(t)) = \max_{\vec{\omega}\in U} H(\hat{T}_*(L,t),\vec{\omega},\vec{p}(t)). \end{cases} \tag{S54}$$

The first equation is called *adjoint equation* and $\vec{p}(t) = (p_1(t),p_2(t)) \in \mathbb{R}^2$ is called *adjoint arc*. The third condition states that the optimal control $u_*(t)$ maximizes the Hamiltonian computed along the optimal trajectory $\hat{T}_*(L,t)$ and the adjoint arc $\vec{p}(t)$. Given $\vec{v} = (v_1,v_2) \in \mathbb{R}^2$, let $\vec{v}^\perp = (-v_2,v_1)$ be the perpendicular vector to $\vec{v}$ obtained by a counterclockwise rotation of $\pi/2$. Define

$$\vec{u}_*(t) := \beta\hat{T}_*^\perp(L,t)\left(\hat{n}\cdot\hat{T}_*^\perp(L,t)\right) \in \mathbb{R}^2 \text{ for all } t\in[0,T_f] \text{ and } \beta \geq 0. \tag{S55}$$

It follows that $\vec{u}_*(\cdot) \in \mathcal{U}$ and the conditions (S54) are satisfied since

$$\begin{cases} \vec{u}_*(t)\cdot\hat{n} = \left(\hat{T}_*^\perp(t)\cdot\hat{n}\right)^2 \geq 0 \\ \vec{p}(t) = 0 \text{ for all } t\in[0,T_f]; \\ H(\hat{T}_*(L,t),\vec{u}_*(t),\vec{p}(t)) = 0 \text{ for all } t\in[0,T_f]. \end{cases} \tag{S56}$$

Finally, let us note that, if $\hat{n}$ is repulsive, the above discussion holds with

$$\vec{u}_*(t) = -\beta\hat{T}_*^\perp(L,t)\left(\hat{n}\cdot\hat{T}_*^\perp(L,t)\right) \in \mathbb{R}^2 \text{ for all } t\in[0,T_f], \beta\geq 0. \tag{S57}$$

It remains to show that the optimal control based approach here proposed agrees with the geometrical description proposed in Bastien et al. (2013) for the gravitational stimulus $\hat{n} = (0,1)$. Replace the optimal solution (equation (S55)) into the dynamics (S50). It follows

$$\begin{cases} \frac{d}{dt}\hat{T}_1(L,t) = -L\hat{T}_1(L,t)\hat{T}_2(L,t); \\ \frac{d}{dt}\hat{T}_2(L,t) = L\hat{T}_1^2(L,t); \end{cases} \tag{S58}$$

Furthermore, the vector $\hat{T}(s,t)$ can be written in terms of the angle $\theta(s,t)$ between $\hat{T}(s,t)$ and the stimulus direction $\hat{n}$:

$$\hat{T}(s,t) = \left(\sin\left(\theta(s,t)\right), \cos\left(\theta(s,t)\right)\right). \tag{S59}$$

Equations (S58) can be expressed as a scalar equation in terms of $\theta(s,t)$, namely

$$\frac{d}{dt}\theta(L,t) = -L\sin\left(\theta(L,t)\right) \tag{S60}$$

and, in view of dynamics (S50), one can write, for any fixed $s \in [0, L]$

$$\frac{d}{dt}\theta(s,t) = -s\sin\left(\theta(L,t)\right). \tag{S61}$$

By differentiating equation (S61) with respect to $s$:

$$\frac{\partial}{\partial t}\frac{\partial}{\partial s}\theta(s,t) = -\sin\left(\theta(L,t)\right) \tag{S62}$$

that is closely related to the dynamics described in Bastien et al. (2013).

#### REFERENCES

Aronna, M. S., Tonon, D., Boccia, A., Campos, C. M., Mazzola, M., Van Nguyen, L., et al. (2017). Optimality conditions (in pontryagin form). In *Optimal Control: Novel Directions and Applications* (Springer). 1–125

Bastien, R., Bohr, T., Moulia, B., and Douady, S. (2013). Unifying model of shoot gravitropism reveals proprioception as a central feature of posture control in plants. *Proceedings of the National Academy of Sciences of the United States of America* 110, 755–760

Silk, W. K. (1989). Growth Rate Patterns which Maintain a Helical Tissue Tube. *J. theor. Biol.* 138, 311–327
